# Cancelling Movement or Pausing Involuntarily? Interpreting Behavioural Delays and Muscle Suppression in Inhibition Tasks

**DOI:** 10.64898/2026.03.03.709399

**Authors:** Simon Weber, Keegan Haugh, Sauro E. Salomoni, Ahreum Lee, Evan J. Livesey, Mark R. Hinder

**Author notes:** Corresponding author: Simon Weber.

## Abstract

A prominent model of action stopping theorises that the reactive cancellation of movement is underpinned by two dissociable processes: a rapid, involuntary “pause” that transiently suppresses motor output, and a slower, voluntary cancellation of movement. As a fundamental component of motor control, the pause process has been posited to generalise broadly to infrequent and salient stimuli (irrespective of whether they bear an imperative to stop) and to be observable as suppression in electromyographical (EMG) recordings.

Over two experiments (N = 24 in each), participants completed standard stop signal and flanker tasks, and novel task variants, where flanking arrows occurred infrequently, with or without a delay (relative to the central go stimulus), or coincident with a stop stimulus.

Relative to a standard flanker task, presenting flankers infrequently specifically increased slowing in incongruent trials after at least three preceding trials without flanking stimuli, with no effect in congruent or neutral trials, indicating involuntary delays to infrequent events are sensitive to both stimulus features and the recency of perceptually similar events.

Presenting flanker arrows of any congruence alongside a stop stimulus resulted in faster EMG suppression than stop stimuli alone, indicating that additional infrequent stimuli can facilitate inhibitory responses. However, presenting flanker stimuli at an equivalent frequency and delay outside of a stopping task did not reliably elicit EMG suppression or behavioural delays. The current findings challenge current models by demonstrating that physiological signatures previously interpreted as manifestations of a generalisable pause process are instead context specific, requiring the anticipation of action cancellation.

## 1. Introduction

Inhibition represents a fundamental aspect of motor control, enabling us to pursue goal-directed behaviours while suppressing inappropriate or erroneous movements (Bari & Robbins, 2013). While the precise subcomponents of inhibition are still a topic of debate (Friedman & Miyake, 2004; Nigg, 2000; Tiego et al., 2018), most contemporary research differentiates between the ability to stop planned or initiated motor responses - usually referred to as motor inhibition - and the ability to suppress responses to distractor information - variably referred to as “interference control”, “perceptual inhibition”, or “attentional inhibition” (Rey-Mermet et al., 2018; Sebastian et al., 2013; Shields & Yonelinas, 2025). Both capacities have been extensively researched in a range of clinical populations (Lipszyc & Schachar, 2010; Rabi et al., 2020), and in relation to healthy aging (Rey-Mermet & Gade, 2018).

Despite the proliferation of research investigating inhibition, there is still uncertainty regarding how we should interpret performance in the behavioural paradigms used to assess it. Motor inhibition is typically assessed with the stop signal task (SST; Verbruggen et al., 2019). These tasks feature a series of trials in which the participant needs to press a button when a visual “go signal” appears. On a subset of trials (usually 25-33%), an auditory or visual “stop signal” occurs after a short delay (referred to as the stop signal delay; SSD), indicating that the participant must *inhibit* their response (i.e., make no button press). Reactively stopping a movement in this manner is thought to rely on a predominantly right-lateralised stopping network connecting frontal areas (pre-supplementary motor area [preSMA], right inferior frontal gyrus [rIFG]) to basal ganglia pathways, which in turn suppresses motor output to the primary motor cortex (M1) to stop movement (Aron et al., 2014; Diesburg & Wessel, 2021). Notably, though, key regions of this stopping network are recruited in other situations, where action cancellation isn’t required, casting doubt on the functional roles they serve (Wessel & Aron, 2017). For example, brain imaging research has demonstrated both activation of the rIFG and behavioural slowing when *initiating* movement, if the stimulus triggering this movement is rendered unexpected by invalid biasing cues (Sebastian et al., 2021). Similarly, a large body of research has found that the rIFG is activated by other infrequent signals presented in the context of a SST, but which do not require stopping (Hampshire et al., 2010; Sharp et al., 2010; Wang, 2025). This has led some authors to posit that SSTs conflate two inhibitory processes – an involuntary suppression that occurs when the stop signal appears, simply on account of it being a salient and unexpected stimulus, and the voluntary cancellation of movement as a result of the imperative to stop (Diesburg & Wessel, 2021).

A recent theoretical model, the “pause then cancel” model, attempts to characterise these two distinct processes, and associate distinct physiological and electrophysiological signatures with each (Diesburg & Wessel, 2021; Schmidt & Berke, 2017). The broadly occurring, involuntary suppression is described as a “pause”, while a slower, voluntary stopping/retuning of movement is “cancel”. Notably, the “pause” has been proposed to inhibit the motor system in response to “any salient event” (Tatz et al., 2024), and to be detectable at the level of the muscle with electromyography (EMG), manifesting as the rapid suppression of initiated muscle activity (Diesburg & Wessel, 2021; Hervault & Wessel, 2025). There exists a large body of work using EMG to characterise stopping processes in action cancellation tasks (Jana et al., 2020; Raud et al., 2022; Salomoni et al., 2025). This is made possible by the fact that, on many successful stop trials (∼40-60%), a movement will have been initiated (the effector muscle is activated), but this is suppressed prior to enough force being generated to yield an overt behavioural response (e.g., a button press), resulting in a “partial response”. By measuring the time from stop signal presentation to the peak muscle activity during the partial response (the point where muscle activity begins to reduce), a single trial measure of stopping speed can be determined at the muscle.

Partial responses have been associated with the pause process based primarily on the observation that muscle activity drops 140-170ms following the stop signal, while signatures purportedly associated with the “cancel” process occur later (e.g., electrophysiological signatures such as the frontocentral P3, occur 200-300ms following the stop signal; Diesburg & Wessel, 2021; Wessel & Aron, 2015). Notably, if partial EMG responses represent a pause process, and this process is reliably triggered in response to any infrequent and salient signals (Tatz et al., 2024), it follows that they should also manifest when salient stimuli are not associated with an imperative to stop. Indeed, there is some evidence of partial responses occurring in response to salient signals that do not require stopping, but these are typically presented *within* SSTs, with similar features, and occurring with a similar frequency, to stop signals (Kemp et al., 2024; Tatz et al., 2024, Weber et al., 2026). As such, the suppression of movement (as a result of the pause) may have little to do with attentional capture associated with an infrequent event, and more to do with the fact that participants must rapidly discriminate between two stimuli, only one of which requires action cancellation (Bissett & Logan, 2014; Sanchez-Carmona et al., 2021; Weber et al., 2026). Limited research has sought to investigate partial responses across both stopping and non-stopping contexts. However, in a recent study, when stopping was not required to any infrequent stimulus (and instead infrequent stimuli required initiating an additional movement), partial responses were much less frequent compared to an SST with perceptually equivalent infrequent stimuli, presented with similar frequency and delays (Weber et al., 2023). Here, we sought to extend our fundamental understanding of how motor control is affected by responses to infrequent salient events by testing specific predictions of the pause then cancel model in a number of novel task contexts.

Partial responses in the context of action cancellation are typically interpreted as inhibitory responses originating at the cortex arriving at the muscle (Raud et al., 2022; Salomoni et al., 2025). However, this same signature has been interpreted differently in different contexts. For example in a choice response task where peripheral or extraneous stimuli introduce conflict between choice options (such as in the Eriksen Flanker task, described below), partial responses have been interpreted as a “change of mind” - i.e., occasions where a response was selected and initiated at the level of the muscle, but abandoned prior to a behavioural response being generated (a button press), and before an alternative response was made instead (Burle et al., 2002; Roger et al., 2014; Servant et al., 2015; Stone et al., 2022).

The sensitivity and signal-to-noise ratio of EMG necessitate that appropriate algorithms are implemented to ensure that the partial responses detected are theoretically plausible (i.e., occur at the correct time relative to the stimulus, and in the expected effector muscle) and do not simply represent sub-threshold EMG that is unrelated to the task. Necessarily, therefore, these algorithms vary across studies according to what the bursts purportedly represent. For example, conceptualising partial responses as a *change of mind*, in a flanker task Servant et al., (2015), filtered out 10.7% of trials, on account of partial responses occurring where they did not match the theoretical interpretation (e.g., if there was a partial response preceding, but on the same side, as the correct response). Similarly, in SSTs, searches for partial responses are normally conducted only in the cued hand, and only *after* presentation of the stop signal (Jana et al., 2020; Raud et al., 2022). No prior research has directly compared the characteristics of partial responses in these contexts, to determine if potential differences in the mechanisms underpinning them (action cancellation/change of mind) manifest at the level of the muscle. Thus, in addition to assessing manifestations of a “pause” response in novel task contexts, we sought to conduct exploratory analyses comparing key physiological parameters (amplitude, offset slope, length, and latency following the signal they relate to) between partial responses associated with a “change of mind” versus those that would typically be interpreted as “action cancellation”.

The Eriksen Flanker Task requires participants to make left-or right-hand responses to a central target (e.g., an arrow or letter) while ignoring surrounding distractor stimuli (i.e. flankers; Eriksen & Schultz, 1979; Ridderinkhof et al., 2021; Xiao et al., 2023). In the case of arrow stimuli, flankers may be congruent (pointing in the same direction as the target), incongruent (pointing opposite to the target) or neutral (neither opposing nor aligning with the target). Congruent flankers facilitate faster reaction times (RTs) relative to neutral and incongruent responses, due to their activation of the same motor response as the target (Ridderinkhof et al., 2021). In contrast, incongruent flankers introduce response conflict, which reliably slows RTs relative to congruent and neutral trials, and increases error rates. Here, we tested whether a rapid “pause” process was involuntarily initiated in a flanker task, by presenting flanker stimuli infrequently (with a frequency that is typical of stop signals in an SST). Electrophysiological data suggests that processing of a stop signal in the rIFG occurs ∼120ms following its presentation (Aron et al., 2014; Jana et al., 2020), and that this is transmitted to M1 (via the basal ganglia) by ∼140ms (Wessel & Aron, 2017), finally resulting in muscle suppression 140-170ms from when the stop signal was presented (Raud et al., 2022; Salomoni et al., 2025). While the effects of flankers are thought to be distributed throughout the sensory hierarchy (Servant et al., 2015), electrophysiological evidence suggests the earliest observable difference between congruent and incongruent trials is an anteriorly distributed negativity measured with scalp EEG ∼200ms post-stimulus (Donohue et al., 2016).

If, as has been proposed, the pause process generalises to other contexts, then presenting flanker stimuli infrequently should also trigger a pause, resulting in slowing (relative to trials where the same peripheral flanker is presented expectedly). Based on the rapidity with which it occurs, we predicted that the pause process would either precede decoding of stimulus features, or be unaffected by them, thus occurring consistently for congruent, incongruent and neutral flanker trials. This prediction is also consistent with the theory, whereby the *salience* of an infrequent event, rather than its association with response options, triggers the pause (Diesburg & Wessel, 2021). To that end, we developed a novel flanker task in which the majority of trials were choice responses with no flanking stimuli, and a subset (equivalent to the typical rate of stop signal presentation in an SST) were equally divided between congruent, incongruent, and neutral flanker trials. Notably, in SSTs, the stop signal occurs *after a delay*, capturing attention at a point when movement is being prepared, or muscle activity has started to manifest. To determine if this delay is required for the pause process to manifest, we also developed a condition in which flankers were presented both infrequently, and after various delays (equivalent to the most common stop signal delays in SSTs).

In sum, the current research firstly sought to assess whether infrequent events trigger a generic “pause” that is impervious to stimulus features in a novel task context in which action stopping is not part of the response set. We predicted that presenting flanker stimuli infrequently would cause a generic slowing for all congruencies (i.e., a main effect of condition when comparing a standard flanker task to one with infrequent stimuli, and no interaction effect with congruency). Secondly, we tested whether presenting flanker stimuli infrequently, *and* after a delay (akin to stop signals in an SST), would cause the pause process to manifest as a partial response, observable with EMG. In the case that it did (as the pause then cancel model would predict) we anticipated that this would occur with the same latency as action cancellation partial responses from the stop signal task (strong Bayesian evidence for the null). Thirdly, we conducted exploratory analyses to compare key characteristics of partial responses associated with “changes of mind” in flanker tasks with those that represented “action cancellation” in the SST, to see if meaningful qualitative differences (on account of muscle suppression being driven by distinct mechanisms) could be observed in EMG profiles.

## 2. Methods

### 2.1. Participants

Participants were recruited through the University of Tasmania’s psychology research participation system, or via direct invitation. Participants were required to have normal or corrected-to-normal vision, and no neurological conditions. For Experiment 1, 26 participants were recruited, though data from 2 participants were excluded on account of psychological states, not disclosed when prompted at the time of consent but later divulged, that may have impacted results (following exclusions: 16 female, 8 male; mean age = 26 years, age SD = 5.80, age range = 18 – 36 years). For Experiment 2, 24 participants were recruited (10 female, 14 male; mean age = 24 years, age SD = 5.86, age range = 18 – 34 years). Participant numbers were determined based on effect size calculations from prior research, where cohorts of this size were sufficient to detect small differences in the latency of EMG activity (Weber et al., 2025). Participants were awarded two hours of research participation credit (a component of their undergraduate study), or a $20 AUD gift voucher. This research was approved by the Tasmanian Human Research Ethics Committee (31750).

### 2.2. Experimental Setup

Participants were seated approximately 80cm from a computer screen with forearms pronated and index fingers resting adjacent to vertically-mounted response buttons, such that abduction of the index fingers would register a button press. Tasks were programmed and run using Psychopy3 (Peirce et al., 2019). Stimuli were presented on a 24-inch monitor (ViewSonic XG2431) with a screen size of 23.8 inches (1920 x 1080 resolution, 240 Hz).

Cambridge Electronic Design’s hardware was used for EMG data acquisition and amplification, and Signal software was used to record the data (Cambridge Electronic Design Ltd.). Two electrodes were arranged in a belly-tendon montage on the first dorsal interosseous (FDI) on each hand, and a grounding electrode was placed on the head of the ulna on each wrist. To determine the correct placement of electrodes, participants performed an isometric abduction of their index finger to activate the FDI, allowing the experimenter to locate positions for one electrode at the tendon insertion at the radial base of the index finger and another at the belly of the muscle.

The EMG signal was observed during a few button presses prior to beginning the experiment, to ensure electrode placement had been optimised. Participants were shown the live trace of EMG (on a nearby monitor, separate to that used to present stimuli) and it was explained that they should try to keep their hands relaxed during the experiment. During the experiment itself this monitor was rotated, such that participants could not see, or be distracted by, the live EMG data, though this remained within the researcher’s view, and if the EMG signal indicated a large amount of background noise they would remind the participant to “please try to keep your hands relaxed between button presses”. EMG activity was recorded on every trial in 2000ms sweeps, beginning 500ms prior to stimulus presentation.

### 2.3. Overview of Experimental Conditions

An overview of the conditions is shown in Figure 1. To address our key research questions without creating an unfeasibly long paradigm, we used a mixed within-and between-subjects design, whereby two distinct experiments were run, and key conditions were common to both. The conditions common to both Experiments 1 and 2 (completed by all participants; N = 48) were the Choice Response Task, the Standard Flanker Task, and the Infrequent Flanker Task. Experiment 1 (N=24) additionally featured the Delayed Infrequent Flanker Task, while Experiment 2 (N = 24) additionally featured the Stop Signal Task, and the Stop With Flankers Task, all of which are described below.

**Figure 1:**
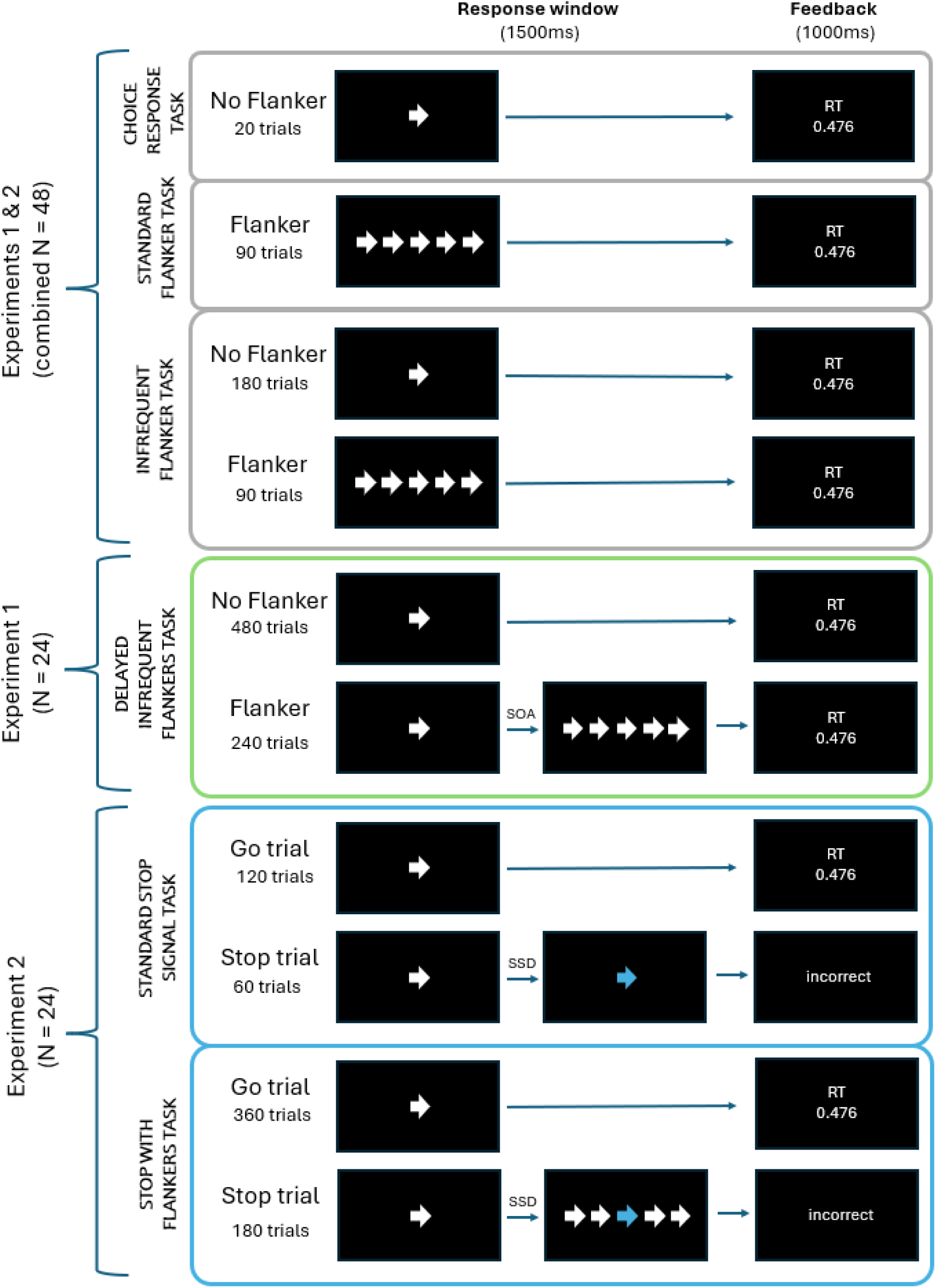
Experiment overview. Three conditions (the Choice Response Task, Standard Flanker Task, and Infrequent Flanker Task) were common to both experiments 1 and 2. Experiment 1 additionally featured the Delayed infrequent Flanker Task, while Experiment 2 additionally featured the Stop Signal Task and Stop With Flankers Task. All trials featuring flanker stimuli were equally divided between congruencies (congruent, incongruent and neutral)

Prior to starting all conditions, participants progressed through comprehensive instruction screens at their own pace. In all conditions, visual stimuli were white (RGB: 255, 255, 255) presented on a black background (RGB: 0, 0, 0), except for stop-signals, which were cyan (RGB: 0, 255, 255). Each trial began with a central fixation cross presented for 1000ms. This was followed by a central arrow pointing either left or right with equal probability (i.e., the target, or go stimulus). Participants were instructed to respond as quickly and accurately as possible to the stimulus with a unimanual button press corresponding to arrow direction, within a 1,500 ms response window. Following each response, performance feedback (described for specific tasks below) was presented in a 1000ms inter-trial interval (ITI) before the next fixation appeared. To minimise fatigue, conditions with more than 90 trials included short rest breaks after every 90 trials (see below for details of trial numbers).

For both experiments, condition order was counterbalanced across participants, though for Experiment 2 we counterbalanced the order of all tasks *barring* the choice response task, which was always conducted immediately prior to whichever stop signal task that participant completed first. This was done because the mean RT from the choice response task was used to calculate the threshold for the proactive slowing warnings (described below).

#### 2.3.1 Choice Response Task

Participants completed 20 choice response trials. Left and right unimanual responses were required to left and right pointing arrows, respectively. Left and right arrows appeared in equal proportion and random order. Mean RT of correct responses was calculated and displayed to participants upon completion.

#### 2.3.2 Standard Flanker Task

Participants completed 90 trials in which the central go stimulus was flanked by two flanker stimuli on either side, presented simultaneously with the central arrow. Participants were required to ignore the direction of the flanking arrows, and respond only to the direction of the central arrow. Flankers could be congruent (pointing in the same direction as the central stimulus), incongruent (pointing in the opposite direction), or neutral (double headed arrows). Notably, not all flanker tasks in the literature utilise neutral trials, and implementation varies in studies that do. Here, we chose to use double-headed arrows (Smith & Ulrich, 2024). Congruent, incongruent and neutral trial types occurred with equal probability (30 of each) and featured an equal number of left and right central stimuli (e.g., 15 left and 15 right per congruency condition). Trials were presented in random order.

#### 2.3.3 Infrequent Flanker Task

This condition was the same as the Standard Flanker Task except that flanker stimuli were only presented on 33% of trials (90 trials of a total of 270 trials), replicating the frequency at which stop signals are presented in SSTs (Verbruggen et al, 2019). The remaining 180 trials were “no-flanker” choice response trials, evenly divided between left and right central stimuli, and identical to the trials used in the Choice Response Task. To ensure that any effects arising from the number of preceding no-flanker trials would not be confounded with congruency effects, we used pseudo-randomised trial orders that controlled the number of (no-flanker) choice response trials preceding each flanker trial. We set the maximum number of no-flanker trials that could occur in a row at four, and for each congruency we ensured there were six trials with each possible number of preceding no-flanker trials (between 0 and 4). Of these six trials, three were left responses, and three were right responses.

#### 2.3.4 Delayed Infrequent Flanker Task

This condition was the same as the Infrequent Flanker Task except that the timing of the infrequent flankers relative to the central target stimulus was manipulated, with flankers appearing at various stimulus onset asynchronies (SOAs; 50, 100, 150, and 200ms) relative to the central target arrow. The total number of trials in this condition was 720, with 240 trials allocated equally to (infrequent) flankers at the four SOAs (20 trials per SOA, per congruency, equal left and right target stimuli). As with the Infrequent Flanker Task, we ensured that any effects from infrequency of flankers would not be confounded with congruency. Accordingly, there were 80 trials per congruency, and within that, 20 per SOA. Thus, there were four trials with each possible number (0-4) of preceding no-flanker trials, with two of these requiring a left response, and two requiring a right response.

#### 2.3.5 Stop Signal Task

This condition featured 180 trials, of which 120 were go trials (67%) and 60 were stop trials (33%). On stop trials, the central arrow changed from white to cyan following a variable stop signal delay (SSD), indicating that participants should cancel their planned or initiated response. If no buttons were pressed, the trial outcome was deemed correct, and the feedback screen read “Correct”; any other response was deemed as a failed stop, and the feedback screen indicated “Incorrect”. SSD started at 150ms and was adjusted by 50ms increments using a staircasing procedure based on stop trial success: successful inhibition made the subsequent stop trial harder by increasing the SSD by 50 ms (up to a maximum of 550 ms), whereas failed inhibition decreased it by 50 ms, to a minimum of 50 ms; (Verbruggen et al., 2019). Trials occurred in pseudorandomised order such that there were never more than two stop trials in a row. Interpreting performance in stop signal tasks can be complicated by the fact that participants tend to slow their responses in anticipation of the stop signals (Langford et al., 2016; Verbruggen et al., 2019). To minimise the degree to which this occurred, we added an additional line of text “you’ve slowed down” to the feedback screen on go trials if RT was >200ms slower than that participant’s median RT in the choice response task (see 2.3.1). In addition to this, the instruction screens for this task told participants to “please try to respond as quickly as possible, trying not to slow down in anticipation of the stop signal”.

#### 2.3.6 Stop With Flankers Task

This condition featured 540 trials. Of these, 67% (360) were go trials, and 33% were stop trials (180). This condition was the same as the Stop Signal Task, except the stop signal (as described above) was coincident with the appearance of flankers. These could be either congruent, incongruent, or neutral (with 60 of each type, equally divided between left and right stops). Separate SSD tracking variables were used for each congruency. Participants were instructed to respond to the go stimulus, and to cancel their response to the go stimulus when the stop stimulus appeared while ignoring the flanker stimuli.

### 2.4 Electromyographical Methods

EMG data were processed with MATLAB (MathWorks, 2018). Initially, each data sweep was filtered with a fourth-order band-pass Butterworth filter at 20-500Hz. Signals were then rectified and low-pass filtered at 50Hz. A single-threshold algorithm was used to detect onset and offset of EMG activity (Hodges & Bui, 1996). A 500ms sliding window was used to establish the period in the 2000 ms sweep with the lowest root mean squared amplitude, which then served as the baseline. Bursts were identified when the EMG amplitude exceeded three SDs above baseline. Bursts with less than 20ms between them were merged into a single event. After burst detection, EMG envelopes were obtained by smoothing the signals at 10Hz. Following this, two types of EMG burst were defined, RT generating bursts, or partial bursts (due to either action cancellation or changes of mind). The RT generating response was defined as the last burst with an onset after the go signal, but before the recorded button press (RT). For a burst to be classed as a partial response its amplitude needed to exceed 10% of the average peak amplitude from that participant’s RT generating responses, from correct trials in that condition (note that in SSTs, this refers to amplitudes from correct go trials - not stop trials, which had no RT generating burst). This was performed separately for each hand. Any partial responses that peaked within 50ms of the go signal appearing were excluded, as these likely reflected activity unrelated to the task (0.21% of partial responses in the non-cued hand and 0.14% of partial responses detected in the cued hand; Jana et al., 2020).

To attain trial-level indices of stopping speed in correct stop trials we subtracted SSD from the latency of the peak of partial response in the cued hand (a measure often refered to as CancelTime) (Raud et al., 2022; Salomoni et al., 2025). Partial responses in the cued hand that peaked < 50ms following the SSD were not included as these likely reflected activity unrelated to the stop signal (Jana et al., 2020; 7.19% of cued hand partial responses in correct stop trials).

Partial responses associated with a change of mind were detected in the non-cued hand in go trials. To ensure that no mirror activation was spurriously interpreted as a change of mind we only included partial responses which peaked prior to the onset of the RT generating response on the cued side.

### 2.5 Data Analysis

#### 2.5.1 Behavioural Data Analysis

Statistical analyses were conducted using generalised linear mixed models (GLMMs) and linear mixed models (LMMs) run in R studio. A model selection approach based on Bayesian information Criteria (BIC) (Schwarz, 1978) was used to determine the optimal structure of random effects.

To assess congruency effects on accuracy in the flanker tasks (categorical data; correct/incorrect) we used binomial model with a probit link function (Caffo & Griswold, 2006). The model included the fixed effects of Condition (Standard Flanker, Infrequent Flanker, Delayed Infrequent Flanker) and Congruence (Congruent, Incongruent, Neutral). The random effects structure included participant intercepts.

For analyses conducted on reaction times, we accomodated for the positive skew (typical of reaction time data) by using models with a gamma distribution and log link function (Lo & Andrews, 2015). To investigate the effects of stimulus infrequency on congruency effects, we ran a GLMM on the data from the Standard Flanker Task and Infrequent Flanker Task. The model included the fixed effects of Congruence (congruent, incongruent, neutral; no-flanker trials in the Infrequent Flanker condition were excluded to avoid issues with missing levels) and Frequency (standard, infrequent). The selected random effects structure included participant intercepts and slopes for Frequency and Congruence.

We predicted that the number of no flanker trials that preceded each flanker trial in the Infrequent Flanker Task would influence the degree of slowing (due to the purported ‘pause’ process) that occurred in the infrequent flanker trials. To investigate this we ran a GLMM on RT of infrequent flanker trials with the fixed effect of Preceding No-Flankers (S, 0, 1, 2, 3, 4), where S represents trials from the Standard Flanker Task, and the other 5 levels represent data from the infrequent flanker task, with the number indicating the count of trials without flankers that preceded that trial. Congruence of the current trial (congruent, incongruent, neutral) was also included as a fixed effect. The selected random effects structure featured participant intercepts and slopes for Preceding No-Flankers.

To investigate the effects of flanker Stimulus Onset Asynchrony (SOA) on RT we ran a GLMM on the data from the Delayed Infrequent Flanker Task, with the fixed effects of Congruence (congruent, incongruent, neutral) and SOA (0ms 50ms, 100ms, 150ms and 200ms), with the latter indicating (in ms) the delay between the central stimulus and the appearance of flanker stimuli. Note that, because all flanker trials in the Delayed Infrequent Flanker Task occurred after some delay (at least 50ms), data from the Infrequent Flanker Task was used for the 0 level of this variable, to enable contrasts (filtered only to include data from participants who completed Experiment 1). The chosen random effects structure included participant intercepts and slopes for SOA.

In SSTs, proactive adjustments may be made to inhibitory networks, to facilitate the reactive stopping that may be required (Lavallee et al., 2014). Behaviourally, this manifests as slowed reaction times (relative to those seen in a task where stopping is not part of the response set) (van de Laar et al., 2011). Here, we assessed this with a GLMM by comparing correct go trials from the Standard SST and the Stop With Flankers tasksto the Choice Response Task. We also included trials without flankers from the Infrequent Flanker and Delayed Infrequent Flanker conditions, to determine whether preparatory adjustments were made when participants anticipated the potential need to suppress *peripheral* stimuli. Thus the single fixed effect was Condition (Choice Response, Standard SST, Stop With Flankers, Infrequent Flanker, Delayed Infrequent Flanker). The random effects structure included only participant intercepts.

The integration method was used to calculate stop signal reaction time (SSRT) in the Standard SST, and seperately for each congruency in the Stop With Flankers condition (Verbruggen et al., 2019). This participant-level data (i.e., one value per condition, for each participant) was analysed using a linear mixed model with participant intercepts. Previous work assessing associations between response inhibition and perceptual inhibition has correlated SSRTs and mean “flanker congruency effects” (Bender et al., 2016; Loomes et al., 2023). We checked for an association in the current experiments with Pearson’s correlations comparing SSRT in the Standard SST to flanker congruency effects in the Standard Flanker Task.

#### 2.5.2 Electromyographical Data Analysis

Two distinct models were run to assess specific questions regarding the *prevalence* of change of mind partial responses. Firstly, we wanted to confirm that changes of mind were more common in flanker trials than non-flanker trials. This was investigated within the infrequent flanker condition (as this featured trials both with and without flankers). A GLMM was run with a binomial distribution (change-of-mind partial response: present or absent) and probit link function with the single fixed factor of Congruence (no_flanker, congruent, incongruent, neutral). Following model selection, the random effects structure featured participant intercepts. A subsequent model (with the same distribution and link function) was conducted to compare the presence/absence of change-of-mind partial respones across the three flanker conditions, with no-flanker trials removed to avoid issues with missing levels in the standard flanker task. The model featured a main effect of Condition (standard, infrequent, delayed) and Congruence (congruent, incongruent, neutral). The selected random effects structure featured participant intercepts.

A model was run to determine if the *latency* of change-of-mind partial responses varied as a result of congruency and flanker frequency. The model featured the fixed effects of Frequency (Standard Flanker, Infrequent Flanker) and Congruence (congruent, incongruent, neutral). As with the above model, no flanker trials from the Infrequent Flanker Task were not included. It was ambiguous from visual inspection of distributions as to whether a Gaussian or gamma distribution should be used, so model selection was conducted to determine which resulted in the lowest BIC. The ultimate model used a gamma distribution and log link function with a random effects structure with participant intercepts and slopes for Frequency.

We also ran a model to assess the latency of action reprogramming in trials featuring a change of mind (specifically, the time from peak of the change of mind partial response to the onset of the RT generating burst). The selected model used a gamma distribution and log link function with a random effects structure featuring participant intercepts.

To test for differences in CancelTime across the stop signal task conditions a GLMM was run with a gamma distribution and log link function and the single fixed factor of Stop Type (standard SST, congruent, incongruent, neutral), with one level coming from the regular SST and the other three levels drawn from the Stop With Flankers Task. The ultimate random effects structure included participant intercepts and slopes for Stop Type. Following visual inspection of CancelTime distributions it was apparent that a pattern of faster stopping in the Stop With Flankers conditions was occurring, but that this was specific to trials at the faster end of the distributions. Currently, there is no overarching consensus for how CancelTime results should be filtered. Most studies include some lower limit, such as the 50ms applied here, and by other groups (Jana et al., 2020), and some employ an upper limit of 1.5X the IQR above the third quartile of the data (Jana et al., 2020). Notably though, other papers report no outlier removal method when determining partial response latency (Raud et al., 2022). With the lack of established conventions in mind, we ran an exploratory follow-up test using more conservative cut-off filters, based on the fact that density plots showed clear differences in peaks between conditions, which were dwarfed by the large positive skew. In this follow-up analysis, we specifically analysed CancelTimes between 50 and 200ms.

Pearson’s correlations compared each participant’s average CancelTime to their SSRT in each of the stopping conditions (neutral and three flanker conditions). We also sought to investigate the effects of number of preceding go trials on stopping efficacy. Including the factor “number of preceding gos” (0, 1, 2, 3, 4) in a model that also included a factor of stop type resulted in issues with model convergence. As such, this was tested in a separate model, in which we pooled data from all stop types (i.e., a single fixed factor of “number of preceding gos”) and which included an ultimate random effects structure of participant intercepts.

Exploratory analyses were conducted to compare a number of key characteristics of partial responses associated with a change of mind (occurring in the non-cued hand in flanker tasks prior to initiation of a correct response) to those associated with action cancellation (occurring in the cued hand following the stop signal in successful stop trials). Each model featured the fixed factors of condition (Standard Flanker, Infrequent Flanker and Stop With Flankers) and congruence (congruent, incongruent, neutral). Models were run on burst amplitude, burst length (onset to offset; defined above), latency (relative to the presentation of flankers, or stop signal, for change of mind and action cancellation, respectively), and offset slope. To enable assessment of amplitude, partial burst EMG profiles were normalised to that participant’s average amplitude of RT generating responses from the choice response task, averaged across all trials (separately for each hand). The ultimate random effects structure for this model featured only participant intercepts. Following visual inspection of distributions and model selection, a gamma distribution and log link function were used for burst length, with a random effects structure of participant intercepts. Following model selection latency was analysed with a gamma distribution and log link function, and a random effects structure of participant intercepts. Offset slope was calculated by determining the amplitude of the (smoothed) EMG envelopes at the time window from the onset to the offsets of the RT-generating burst. Then, the derivative at each sample was determined using the MATLAB function ‘diff’ (to determine the difference between consecutive samples). The point of minimum (most negative) value within the defined range of the partial burst was identified, and normalised to the peak amplitude of the partial response. The selected model for slope (rate of offset of the burst) used a Gaussian distribution, identity link function, and participant intercepts in the random effects structure.

## 3. Results

### 3.1 Behavioural results

#### 3.1.1 Flanker Condition Accuracy

Mean percentage of correct responses for each condition and congruence are presented in Table 1. The model revealed a main effect of congruence, χ^2^(2) = 101.34, *p* < .001, a main effect of condition, χ^2^(2) = 11.71, *p* = .002, and a significant interaction effect, χ^2^(2) = 49.54, *p* < .001. Holm-corrected post-hoc tests revealed signficant effects of congruency in the Standard Flanker Task, whereby accuracy on incongruent trials was significantly lower than for congruent trials (*z* = 5.74, *p* < 0.001), and neutral trials (*z* = 4.62, *p* < 0.001), though no significant difference was observed between neutral and congruent trials (*z* = 1.67, *p* = 0.094). The same pattern was observed in the Infrequent Flanker Task (congruent-incongruent, *z* = 8.16, *p* < 0.001; neutral-incongruent, *z* = 7.01, *p* < 0.001; congruent-neutral, *z* = 1.91, *p* = 0.056). In contrast, no significant congruency effects were found in the delayed infrequent flanker condition (congruent-incongruent, *z* = 0.45, *p* = 1.000; neutral-incongruent, *z* = 0.98, *p* = 0.993; congruent-neutral, *z* = 0.53, *p* = 1.000).

**Table 1:** Behavioural Results (RT and accuracy) from Flanker Tasks.

|  | No Flanker | Congruent | Incongruent | Neutral |
| --- | --- | --- | --- | --- |
| **Choice Response** |  |  |  |  |
| Accuracy (%) | 99.17% | - | - | - |
| RT (ms) [95%CI] | 383 [376-391] | - | - | - |
| **Standard Flanker** |  |  |  |  |
| Accuracy (%) | - | 99.31% | 95.76% | 98.68% |
| RT (ms) [95%CI] | - | 396 [374-420] | 437 [414-461] | 405 [382-428] |
| **Infrequent Flanker** |  |  |  |  |
| Accuracy (%) | 98.97% | 98.82% | 92.15% | 97.92% |
| RT (ms) [95%CI] | 384 [379-389] | 390 [372-410] | 462 [441-483] | 409 [389-429] |
| **Delayed Infrequent Flanker** |  |  |  |  |
| Accuracy (%) | 97.92% | 97.35% | 97.10% | 97.61% |
| RT (ms) [95%CI] | 372 [367-391] | 375 [358-397] | 398 [375-421] | 384 [362-407] |

RT data represents estimated marginal means (and associated 95% confidence intervals, CI) from associated GLMMs. Accuracy data represents descriptive data.

#### 3.1.2 Flanker Conditions Reaction Times

Results from the model comparing flanker trials in the Standard Flanker Task and Infrequent Flanker Task are depicted in Figure 2, and estimated marginal means are represented in Table 1. The model revealed no significant main effect of frequency, χ^2^(1) = 0.581, *p* = 0.446, though there was a main effect of congruence, χ^2^(2) = 227.93, *p* < 0.001, and a significant interaction effect, χ^2^(1) = 65.55, *p* < 0.001. Holm-corrected post-hoc tests investigating the main effect of congruence revealed congruent trials were faster than incongruent trials (*z* = 14.46, *p* < 0.001, *d* = 0.54), and neutral trials, (*z* = 4.57, *p* < 0.001, *d* = 0.14), and neutral trials were faster than incongruent trials (*z* = 12.64, *p* < 0.001, *d* = 0.41). The interaction effect was investigated with holm-corrected post-hoc tests comparing infrequent and standard conditions at each level of congruency. This revealed that presenting flankers infrequently (compared to on every trial) slowed RTs in incongruent trials, (*z* = 2.462, *p* = 0.014, *d* = 0.22), though had no significant influence on RT in congruent, (*z* = 0.674, *p* = 0.500, *d* = 0.06), or neutral, (*z* = 0.44, *p* = 0.664, *d* = 0.04), trials.

**Figure 2:**
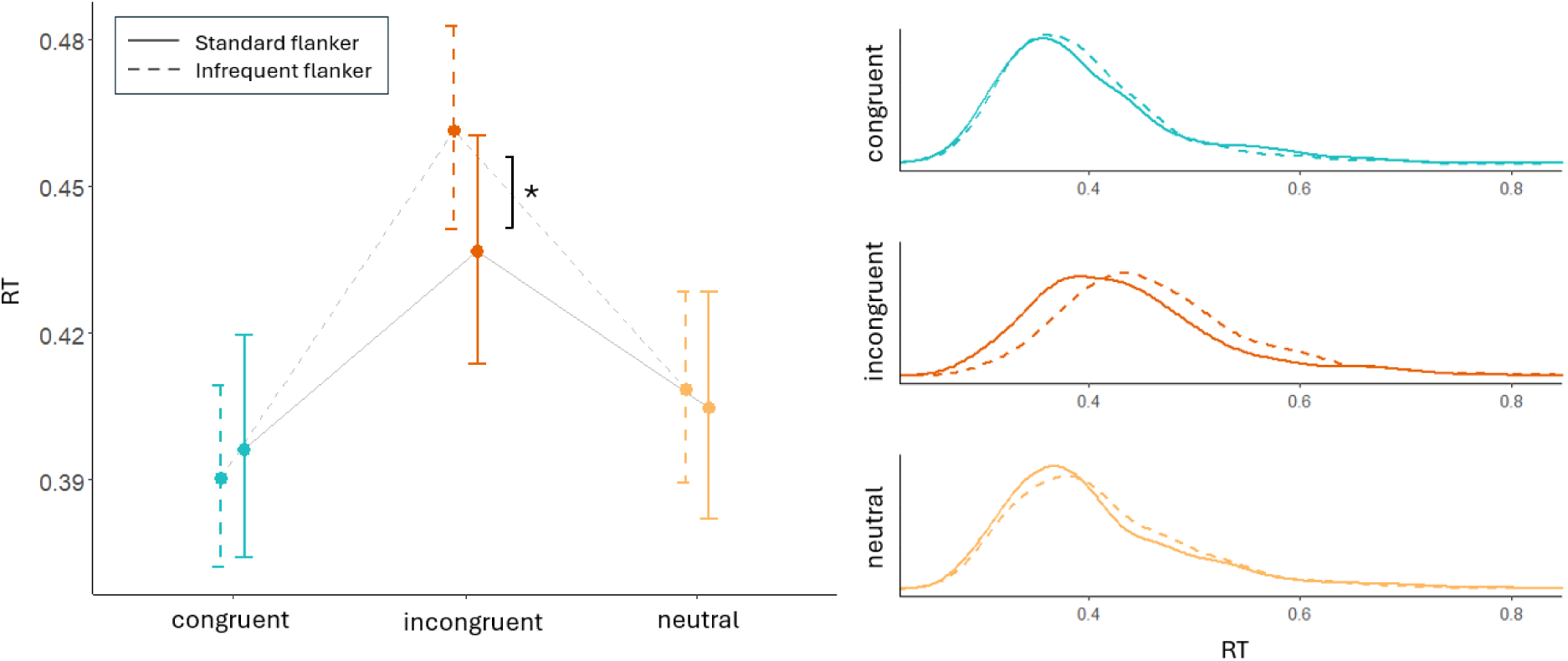
Comparisons of RT in standard and infrequent flanker conditions at each level of congruency revealed presenting flankers infrequently exclusively slowed responses when trials were incongruent. RT distributions for each conguence (disinct lines for the two conditions) are represented in the right panel. * p = 0.014.

The model investigating any effects on RT related to the number of preceding trials without flanker stimuli in the Infrequent Flanker Task revealed significant main effects of congruence, χ^2^(1) = 907.27, *p* < 0.001 and number of preceding gos, χ^2^(1) = 34.70, *p* < 0.001, as well as a significant interaction effect χ^2^(1) = 113.50, *p* < 0.001 (figure 3a). Holm-corrected posthoc tests compared each level of preceding gos (0-4) to the trials from the Standard Flanker Task for each congruence separately. In the neutral and congruent trials, RTs in all levels of preceding gos (0-4) did not vary significantly relative to the Standard Flanker Task (all *p* > 0.44, all *z* < 1.7). However, for incongruent trials, while there was no effect of flanker infrequency with 0 preceding gos (*z* = 0.96, *p* = 0.672, *d* = 0.07), 1 preceding go (*z* = 0.94, *p* = 0.672, *d* = 0.09) 2 preceding gos (*z* = 1.78, *p* = 0.227, *d* = 0.18), or 3 preceding gos (*z* = 2.15, *p* = 0.128, *d* = 0.21), there was a significant effect after 4 preceding gos (*z* = 4.71, *p* < 0.001, *d* = 0.47).

**Figure 3:**
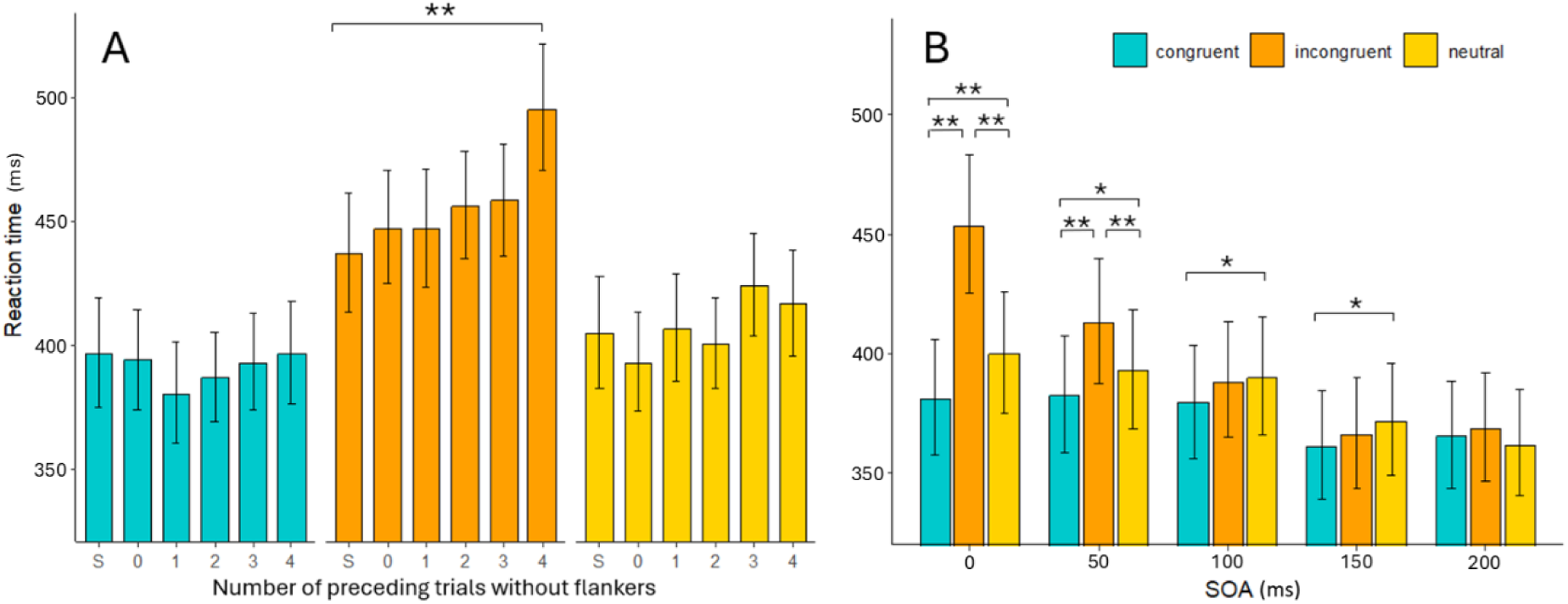
Results from GLMMs investigating effects of flanker infrequency and latency. **A)** “S” represents data from the standard flanker task, used as a baseline to compare to the infrequent flanker task. Significant slowing compared to the standard task occurred only for incongruent flankers after 4 trials without flankers preceded it. **B)** Comparisons of congruency effects at each level of flanker stimulus onset asynchrony (SOA) in the delayed infrequent flanker task. Note that SOA “0” represents data from the infrequent flanker task. ** = p < 0.001 * = p < 0.05.

The results from the model assessing the effect of flanker onset delay (SOA) on RT in the delayed infrequent flanker condition revealed significant main effects of congruence, χ^2^(1) = 153.35, *p* < 0.001, and SOA, χ^2^(1) = 85.18, *p* < 0.001, alongside a significant interaction effect χ^2^(1) = 232.10, *p* < 0.001 (figure 3b). Holm-corrected post-hoc tests checked for effects of congruence at each SOA. At an SOA of 0s, all comparisons were statistically significant (congruent-incongruent, *z* = 18.85, *p* < 0.001, *d* = 0.78; congruent-neutral, *z* = 5.24, *p* < 0.001, *d* = 0.21; incongruent-neutral, *z* = 13.71, *p* <0.001, *d* = 0.57). At an SOA of 0.05s, all comparisons remained significant (congruent-incongruent, *z* = 6.87, *p* < 0.001, *d* = 0.29; congruent-neutral, *z* = 2.45, *p* = 0.014, *d* = 0.13; incongruent-neutral, *z* = 4.43, *p* <0.001, *d* = 0.19). At an SOA of 0.10s, the expected pattern of congruency was only partially evident (congruent-incongruent, *z* = 2.13, *p* = 0.066, *d* = 0.11; congruent-neutral, *z* = 2.51, *p* = 0.036, *d* = 0.13; incongruent-neutral, *z* = 0.39, *p* = 0.698, *d* = 0.02). A similar result was observed at an SOA of 0.15s (congruent-incongruent, *z* = 1.22, *p* = 0.286, *d* = 0.06; congruent-neutral, *z* = 2.69, *p* = 0.022, *d* = 0.13; incongruent-neutral, *z* = 1.47, *p* = 0.286, *d* = 0.06). No congruency effects were significant at an SOA of 0.2s (congruent-incongruent, *z* = 0.80, *p* = 0.761, *d* = 0.04; congruent-neutral, *z* = 0.88, *p* = 0.761, *d* = 0.04; incongruent-neutral, *z* = 1.68, *p* = 0.278, *d* = 0.08).

#### 3.1.3 Proactive Slowing

RT distributions for go trials are depicted in Figure 4. The model run to assess proactive slowing revealed a main effect of condition χ^2^(4) = 2305.12, *p* < 0.001. Holm corrected post-hoc tests revealed significant proactive slowing in the Standard SST (*z* = 23.61, *p* < 0.001, *d* = 0.67), and the Stop With Flankers Task (*z* = 21.18, *p* < 0.001, *d* = 0.56) relative to the choice reaction time task. Comparison of the two SSTs demonstrated that proactive slowing was greater in the Standard SST than the Stop With Flankers Task (*z* = 7.47, *p* < 0.001, *d =* 0.12). While RTs were not significantly different between the Choice Response task and the no-flanker trials in the Infrequent Flanker Task (*z* = 0.20, *p* < 0.851, *d* = 0.01), there was a *speeding* of responses in no-flanker trials in the Delayed Infrequent Flanker Task (*z* = 3.83, *p* < 0.001, *d* = 0.12).

**Figure 4:**
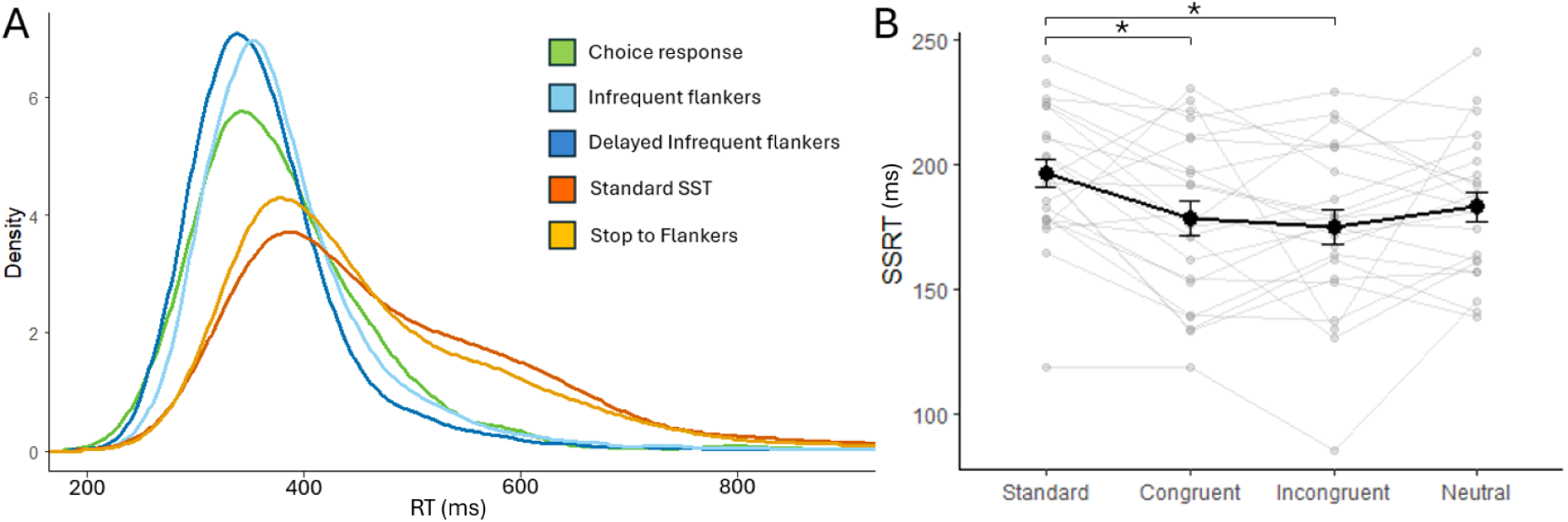
The left panel depicts distributions of reaction times from trials with neither flankers nor stop signals. The significant proactive slowing that occurred in the stop stop signal conditions (Standard SST and Stop With Flankers) is clearly visible. The right panel reflects SSRTs from the standard SST, and the three types of stop trial in the Stop With Flankers condition. Greyed data represents participant-level results, dark line and 95% error bars are group average data. * = p < 0.01.

#### 3.1.4 SSRT

Valid calculation of SSRT requires that accuracy on stop trials approximates to 50% and that failed stop RTs are faster than go RTs (Verbruggen et al., 2019). Both of these requirements were met at the group level (Table 3), and for each individual participant for each condition/congruency. The lowest accuracy rating was 47.06%, and the highest was 63.3%, both of which are still within recommended ranges for SSRT calculation via the integration method (Verbruggen et al., 2019). One SSRT value was removed in one condition on account of it being too fast to be a valid representation of stopping speed (50ms in the neutral condition). Mean SSRTs are represented in Table 2 and Figure 4B. The LMM run on this participant-level data revealed a main effect of condition *F*(3,66) = 6.09, *p* = 0.001. Holm-corrected post-hoc tests revealed SSRT to be longer in the standard SST than stopping with congruent, *t*(23) = 3.33, *p* = 0.007, *d* = 0.59, or incongruent flankers *t*(23) = 3.99, *p* = 0.001, *d* = 0.70 from the Stop With Flankers condition, although the comparison with neutral trials did not reach significance *t*(23) = 2.47, *p* = 0.065, *d* = 0.48. Within the Stop With Flankers condition, comparisons were not significant between congruent and incongruent *t*(23) = 0.66, *p* = 0.785, *d* = 0.11, congruent and neutral, *t*(23) = 0.86, *p* = 0.785, *d* = 0.15, or incongruent and neutral *t*(23) = 1.52, *p* = 0.399, *d* = 0.26. The correlation run on SSRT and flanker congruency effects calculated in the standard flanker condition revealed no significant association between behavioural indices of motor inhibition (SSRT) and perceptual inhibition (flanker congruency effects) *r* = -0.051, *p* = 0.812.

**Table 2:** Estimated marginal means from additional RT models.

| Infrequent Flanker RTs split by number of preceding go trials |  |  |  |  |  |  |
| --- | --- | --- | --- | --- | --- | --- |
|  | Standard Flanker Task | 0 | 1 | 2 | 3 | 4 |
| **Congruent** | 396 [375-419] | 394 [374-414] | 380 [360-401] | 387 [369-405] | 393 [374-413] | 396 [376-417] |
| **Incongruent** | 437 [413-462] | 447 [425-471] | 447 [423-471] | 456 [435-478] | 458 [436-482] | 496 [470-522] |
| **Neutral** | 405 [383-428] | 393 [373-413] | 407 [386-429] | 400 [382-419] | 424 [403-445] | 417 [396-439] |
| Delayed Infrequent Flanker RTs split by Flanker onset asynchrony |  |  |  |  |  |  |
|  | 0 | 0.05 | 0.1 | 0.15 | 0.20 |  |
| **Congruent** | 381 [358-406] | 382 [359-407] | 379 [356-404] | 361 [339-385] | 365 [344-388] |  |
| **Incongruent** | 453 [425-483] | 413 [387-440] | 388 [365-413] | 366 [344-390] | 369 [347-392] |  |
| **Neutral** | 400 [375-426] | 393 [369-419] | 390 [366-415] | 372 [350-396] | 362 [341-385] |  |

RT data represents estimated marginal means (and associated 95% confidence intervals, CI) from associated GLMMs

**Table 3:**
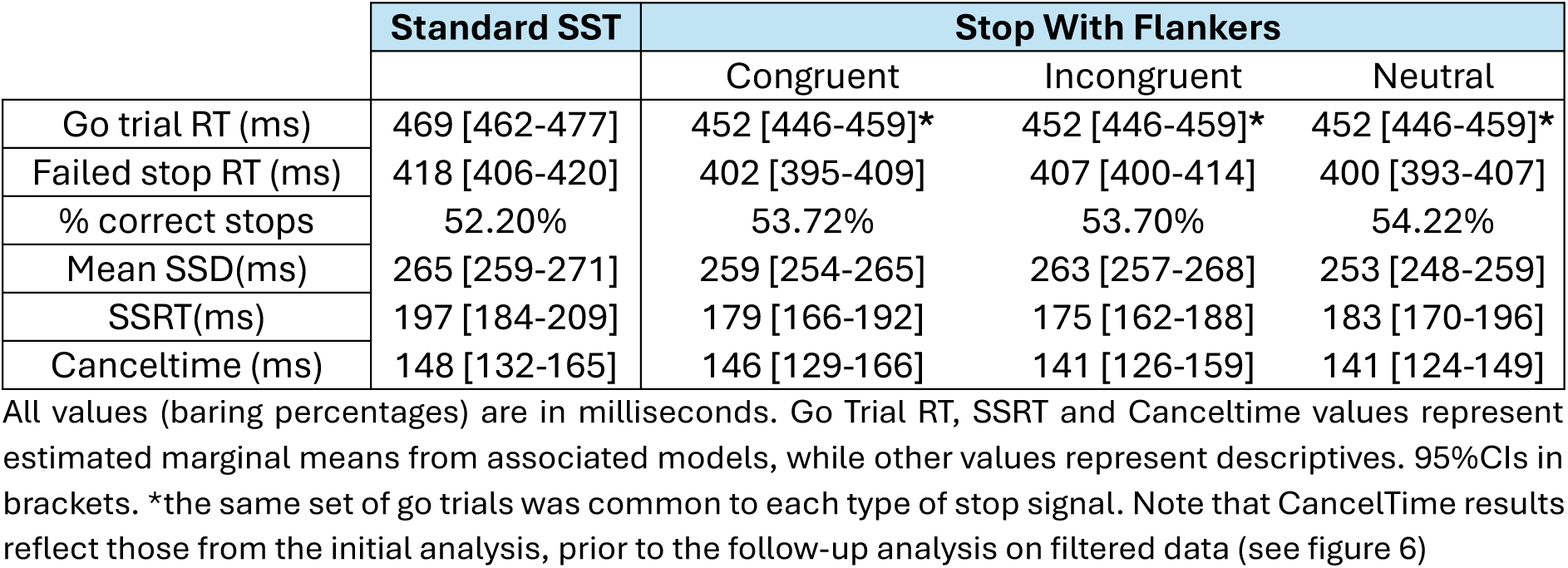
Key results from stop signal conditions.

|  | Standard SST | Stop With Flankers |  |  |
| --- | --- | --- | --- | --- |
|  |  | Congruent | Incongruent | Neutral |
| Go trial RT (ms) | 469 [462-477] | 452 [446-459]* | 452 [446-459]* | 452 [446-459]* |
| Failed stop RT (ms) | 418 [406-420] | 402 [395-409] | 407 [400-414] | 400 [393-407] |
| % correct stops | 52.20% | 53.72% | 53.70% | 54.22% |
| Mean SSD(ms) | 265 [259-271] | 259 [254-265] | 263 [257-268] | 253 [248-259] |
| SSRT(ms) | 197 [184-209] | 179 [166-192] | 175 [162-188] | 183 [170-196] |
| Canceltime (ms) | 148 [132-165] | 146 [129-166] | 141 [126-159] | 141 [124-149] |

All values (barring percentages) are in milliseconds. Go Trial RT, SSRT and Canceltime values represent estimated marginal means from associated models, while other values represent descriptives. 95% CIs in brackets. \*the same set of go trials was common to each type of stop signal. Note that CancelTime results reflect those from the initial analysis, prior to the follow-up analysis on filtered data (see figure 6)

### 3.2 Electrophysiological results

#### 3.2.1 Prevalence of Partial Responses in Flanker Conditions

The percentage of correct response trials in which partial responses were manifested prior to the RT-generating burst, in both the cued hand (the hand on the side indicated by the go stimulus) or non-cued hand, for each flanker condition, and for each congruency, are represented in Table 4.

**Table 4:** Prevalence of Partial Responses in Flanker conditions

| **Flanker tasks – non-cued hand – go trials “changes of mind”** |  |  |  |  |
| --- | --- | --- | --- | --- |
|  | no flanker | congruent | incongruent | neutral |
| Choice response task | 13.45% | - | - | - |
| Standard Flanker task | - | 9.30% | 20.16% | 11.40% |
| Infrequent Flanker task | 10.48% | 11.45% | 29.39% | 13.62% |
| Delayed infrequent Flanker task | 15.72% | 15.98% | 17.21% | 17.48% |
| **Flanker tasks – cued hand – go trials** |  |  |  |  |
| Choice response task | 5.25% | - | - | - |
| Standard Flanker task | - | 5.17% | 8.27% | 4.86% |
| Infrequent Flanker task | 3.80% | 5.34% | 12.21% | 7.66% |
| Delayed infrequent Flanker task | 4.17% | 4.85% | 7.51% | 6.27% |

The results from the GLMM comparing prevalence of change of mind partial responses across conditions (see Figure 5a) revealed a main effect of congruence χ^2^(2) = 197.69, *p* < 0.001, a main effect of condition χ^2^(2) = 29.44, *p* < 0.001, and a significant interaction effect χ^2^(4) = 102.52, *p* < 0.001. Holm corrected post-hoc tests were run testing the effects of condition in each level of congruence. Compared to the Standard Flanker Task there were significantly more changes of mind to congruent trials in the Delayed Infrequent Flanker Task than the Standard Flanker Task (*z* = 4.74, *p* < 0.001), and the Infrequent Flanker Task (*z* = 2.12, *p* = 0.034), and significantly more changes of mind in the delayed than the Delayed Infrequent Flanker Task than the Infrequent Flanker Task (*z* = 2.58, *p* = 0.019). In incongruent trials, there were significantly more changes of mind in the Infrequent Flanker Task than the Standard Flanker Task (*z* = 5.70, *p* < 0.001) and the Delayed Infrequent Flanker Task (*z* = 9.33, *p* < 0.001), and significantly more changes of mind in the Infrequent Flanker Task than the Delayed Infrequent Flanker Task (*z* = 9.33, *p* < 0.001). In neutral trials there were significantly more changes of mind in the Delayed Infrequent Flanker Task than the Standard Flanker Task (*z* = 3.81, *p* < 0.001), though no significant difference between the Infrequent Flanker Task and the Standard Flanker Task (*z* = 2.00, *p* = 0.092) or the Delayed Infrequent Flanker Task (*z* = 1.75, *p* = 0.092).

**Figure 5:**
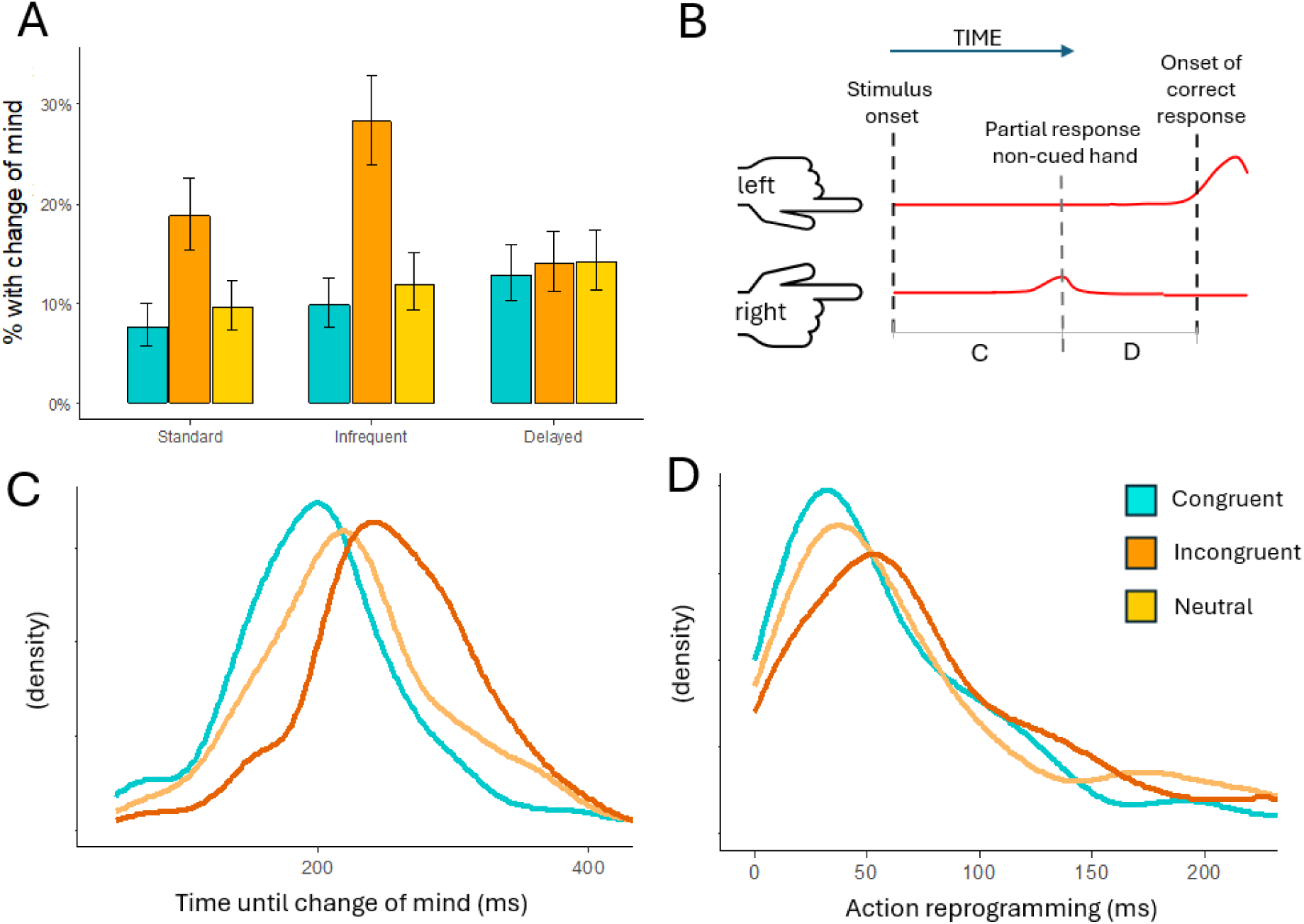
changes of mind in flanker conditions. **A)** Results of the GLMM assessing prevalence of change of mind partial responses in each condition. **B)** Schematic representing smoothed EMG activity. A change of mind partial response involves activation on the non-cued side (in this example, the right hand) which does not reach threshold, followed by a correct response on the cued side. **C)** Density distributions of the time from stimulus onset until the peak of change of mind partial responses. Results are pooled across standard and infrequent flanker tasks, following no main effect of condition. All comparisons between congruencies were statistically significant. **D)** Distributions of the time from change of mind partial responses until EMG onset of the RT generating response (action reprogramming). These distributions are specifically drawn from the Infrequent Flanker task where congruent and incongruent trials were significantly faster and slower than neutral trials, respectively (noting there were no significant congruency effects in the Standard Flanker Task). Note that values less than zero were excluded based on the possibility that such trials may represent mirror activity.

#### 3.2.2 Latency of change of mind partial responses

The model run on latency of change of mind partial bursts revealed a significant main effect of congruence, χ^2^(2) = 99.89, *p* < 0.001, whereas the main effect of condition (standard or infrequent) χ^2^(1) = 0.58, *p* = 0.446, and the interaction effect χ^2^(2) = 1.75, *p* = 0.418, were not significant. Holm-corrected Post-hoc tests run on congruence revealed changes of mind were faster in congruent than both incongruent trials (*z* = 9.57, *p* < 0.001, *d =* 0.63), and neutral trials (*z* = 3.53, *p* < 0.001, *d* = 0.22). Changes of mind were also faster in neutral trials than incongruent trials (*z* = 6.07, *p* < 0.001, *d* = 0.39). Estimated marginal means for the latency of changes of mind in congruent, incongruent and neutral trials (95%CIs in brackets) were 220ms [206-234ms], 268ms [253-284ms] and 238ms [224-253ms], respectively. Density pots of this data are depicted in model 5c.

#### 3.2.3 Action reprogramming after change of mind

The model checking for differences in action reprogramming speed in flanker tasks revealed a significant main effect of congruence, χ^2^(2) = 10.0, *p* = 0.007, a main effect of condition χ^2^(1) = 7.17, *p* = 0.007, and a signfiicant interaction, χ^2^(1) = 8.62, *p* = 0.013. Holm corrected post-hoc tests revealed that there were no significant effects of congruence in the Standard Flanker Task when comparing congruent and incongruent (*z* = 0.17, *p* = 0.867, *d =* 0.013), congruent and neutral (*z* = 1.71, *p* = 0.173, *d* = 0.18) and incongruent and neutral (*z* = 2.18, *p* = 0.08, *d* = 0.18). Estimated marginal means for reproragmming times in the Standard Flanker Task for congruent, incongruent and neutral trials (95%CIs in brackets) were, 74ms [62-89ms], 75ms [65-87ms] and 62ms [53-74ms], respectively.

In the Infrequent Flanker Task reprogramming was faster in congruent trials than incongruent trials (*z* = 3.808, *p* < 0.001, *d* = 0.32) and neutral trials (*z* = 2.27, *p* = 0.046, *d* = 0.21). There was no significant difference between incongruent and neutral trials (*z* =1.30, *p* = 0.193, *d =* 0.10). Estimated marginal means for reproragmming times in the Infreqent Flanker Task for congruent, incongruent and neutral trials (95%CIs in brackets) were, 68ms [57-80ms], 93ms [82-106ms] and 84ms [72-99ms], respectively. Density plots for action reprogramming in this condition are depicted in Figure 5d.

#### 3.2.4 Prevalence of partial responses in stopping tasks

The percentage of correct stop trials with partial responses, and – as a comparison – the number of partial bursts that occurred with the same constraints in the Delayed Infrequent Flanker Task are depicted in Table 5. The constraints (SSD +50ms; SOA +50ms) represent those which were used for the CancelTime analysis (and the equivalent constraints that would be used to calculate an equivalent measure in the Delayed Infrequent Flankers Task). As seen in table 5, the prevalence of partial responses in stop trials is high (>55% of successful stop trials in all conditions/congruencies) Despite the large number of delayed infrequent flanker trials, there were very few partial responses observed in a commensurate time window (insufficient to consider these signatures a robust measure - 3.30%, 6.02% and 3.93% of correct trials in each congruency, respectively).

**Table 5:** Prevalence of Partial Responses in the Stop Signal Tasks and Delayed Infrequent Flanker Task.

| **Stop signal tasks - cued hand after SSD+50ms (for CancelTime)** |  |  |  |  |
| --- | --- | --- | --- | --- |
|  | **no flanker** | **congruent** | **incongruent** | **neutral** |
| Standard Stop Signal Task | 60.36% | - | - | - |
| Stop With Flankers Task | - | 55.89% | 56.04% | 56.56% |
| **Delayed Flankers - cued hand after SOA+50ms** |  |  |  |  |
| Delayed Infrequent Flankers Task | - | 3.30% | 6.02% | 3.93% |

*Values represent percentage of correct trials.*

#### 3.2.5 CancelTime

The model to determine whether the number of go trials that preceded a stop signal affected physiological stopping speed revealed no significant main effect χ^2^(7) = 7.20, *p* = 0.126. A follow-up Bayesian ANOVA revealed extreme evidence for the null (BF_01 =_680.44). Estimated marginal means for each number of preceding gos (with 95%CIs in brackets) were as follows: 0 = 141 ms [127-157], 1 = 145 ms [130-161], 2 = 146 ms [131-163], 3 = 147 ms [132-164], 4 = 152 ms [136-169]. This was assessed in a separate model to that which compared CancelTimes in different conditions, due to model convergence issues (see section 2.5.2).

The model comparing CancelTime between stop signal types revealed no significant main effect χ^2^(3) = 2.79, *p* = 0.426. Estimated marginal means are depicted in Table 3 and CancelTime distributions are presented in Figure 6. Re-running the CancelTime analysis with all CancelTimes that were slower than 200ms excluded (as per Methods) revealed a significant main effect of stop type χ^2^(3) = 56.46, *p* < 0.001. Holm-corrected post-hoc tests revealed stopping in the standard SST to be significantly slower than stops with congruent (*z* = 6.23, *p* <0.001, *d* = 0.43), incongruent, (*z* = 6.23, *p* <0.001, *d* = 0.39), and neutral stop signals (*z* = 6.23, *p* <0.001, *d* = 0.43). No significant differences in stopping speed between the stops with flankers were observed. Estimated marginal means were 130ms in the standard stopping task and 117ms, 118ms, and 117ms in congruent, incongruent, and neutral stop signals with flankers, respectively.

**Figure 6:**
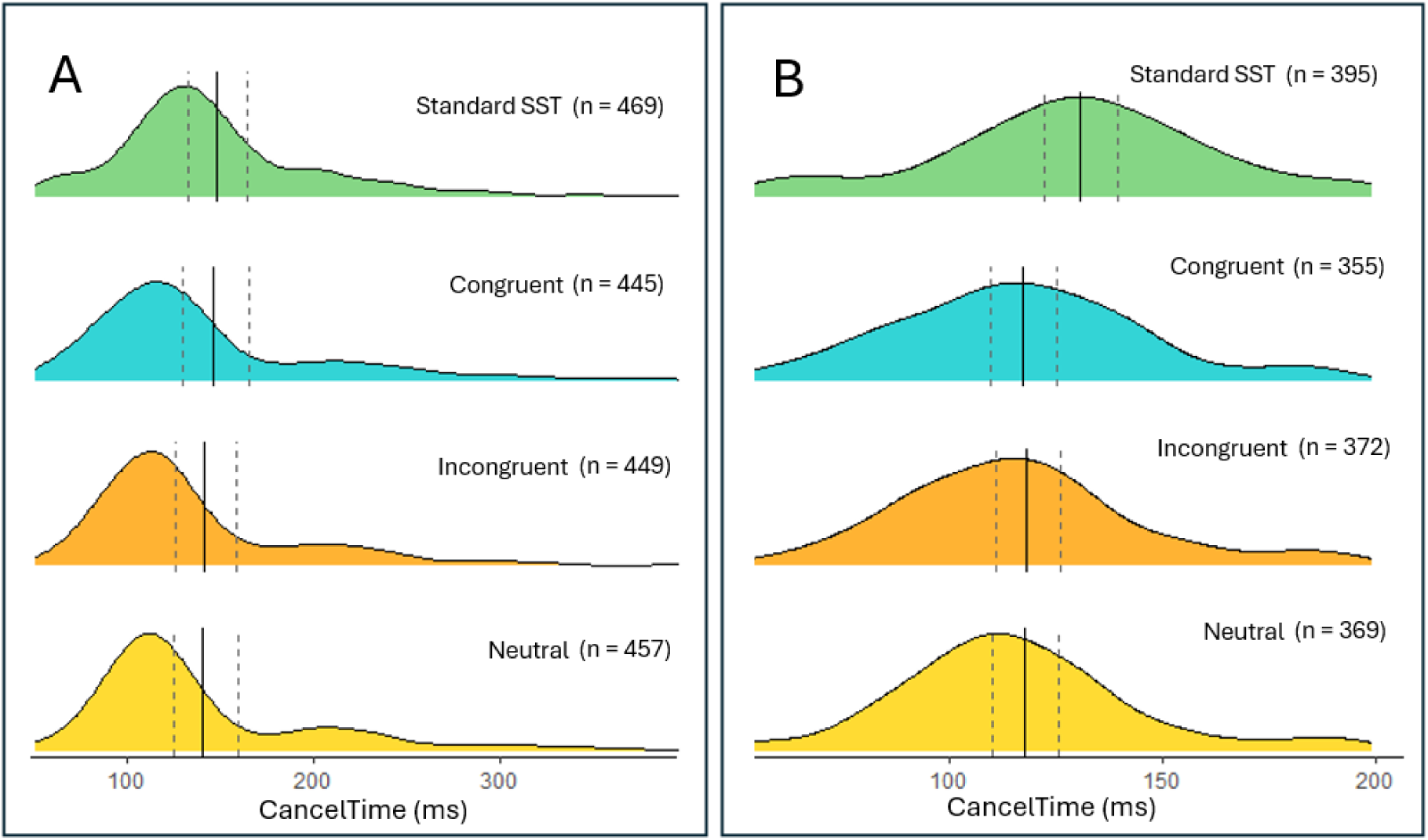
Canceltime distributions from the standard SST condition, and each level of congruence from the Stop With Flankers condition. **A)** Distributions with no upper threshold. Solid lines represent estimated marginal means, while dashed lines represent lower and upper 95%Cis, exhibiting the right-shifting effect of long CancelTimes on mean values. **B)** Distributions with rigorous upper limits (Canceltime <200ms) applied, as per our exploratory analysis.

Using the initial CancelTime analysis, significant possitive correlations were found between participant mean CancelTimes and SSRTs for congruent stops (*r* = 0.78, *p* = <0.001) and neutral stops (*r* = 0.53, *p* = 0.009), but not for stops from the standard SST (*r* = 0.33, *p* = 0.122) or the incongruent stops (*r* = 0.31, *p* = 0.145). Correlation plots can be found in the supplementary materials.

#### 3.2.6 Comparison of change of mind and action cancellation partial responses

The results of the analyses comparing partial reponses representing changes of mind and those associated with action cancellation are depicted in Figure 7, with estimated marginal means for each model presented in Table 6. Amplitude units are relative to the amplitude of RT generating responses in the Choice Response Task (see section 2.5.2 for details regarding each measure).

**Figure 7:**
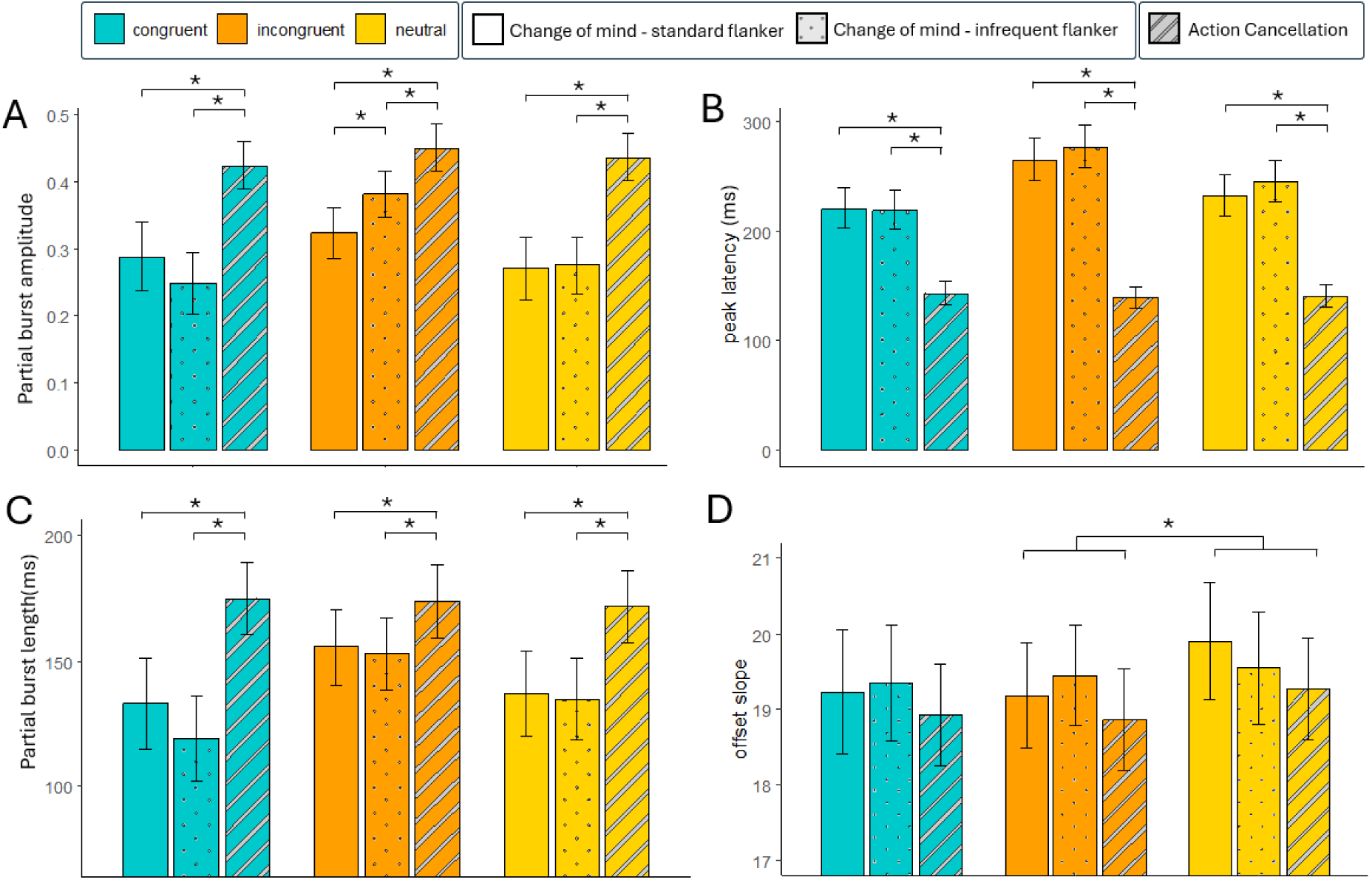
Each plot represents a different partial response attribute divided into trial congruence (by colour) and condition (by pattern) **A)** amplitude of partiial burst peaks. **B)** the latency of the peak of the burst relative to the signal it relates to. For change of mind partials this is the appearance of the flanking stimuli, while for action cancellation partials this represents the appearance of the stop signal. **C)** the length of the partial burst (from offset to onset)**. D)** the steepness of the offset (offset slope) of partial responses, following their peak. Greater values represent steeper slopes. There was a significant difference between incongruent and neutral trials, but no interaction with condition. **\*** = p <0.05.

**Table 6.** Partial Response Attributes for Change of Mind *versus* Action Cancellation. Brackets indicate 95%CIs from models.

|  |  | Change of Mind |  | Action Cancellation |
| --- | --- | --- | --- | --- |
|  |  | Standard Flanker | Infrequent Flanker | Stop With Flankers |
| **Amplitude** | *Cong* | 0.289 [0.238-0.339] | 0.247 [0.201-0.293] | 0.424 [0.389-0.458] |
|  | *Incong* | 0.323 [0.285-0.361] | 0.382 [0.347-0.416] | 0.450 [0.416-0.485] |
|  | *neut* | 0.271 [0.224-0.317] | 0.275 [0.232-0.318] | 0.436 [0.402-0.471] |
| **Peak Latency (ms)** | *Cong* | 221 [203-240] | 219 [202-238] | 144 [134-154] |
|  | *Incong* | 265 [246-285] | 277 [258-297] | 139 [130-150] |
|  | *neut* | 233 [215-252] | 245 [227-265] | 141 [131-151] |
| **Length (ms)** | *Cong* | 133 [115-151] | 119 [102-136] | 175 [160-189] |
|  | *Incong* | 156 [140-171] | 153 [139-168] | 174 [159-188] |
|  | *neut* | 137 [120-154] | 135 [119-151] | 172 [157-186] |
| **Offset Slope** | *Cong* | 19.2 [18.6-20.1] | 19.3 [18.6-20.1] | 18.9 [18.3-19.6] |
|  | *Incong* | 19.2 [18.5-19.9] | 19.5 [18.8-20.1] | 18.9 [18.2-19.5] |
|  | *neut* | 19.6 [18.8-20.2] | 19.9 [19.1-20.7] | 19.3 [18.6-19.9] |

The models for amplitude, length and latency all revealed significant main effects of congruence and condition, and a significant interaction effect. For conciseness, we summarise findings here, with full results reported in the supplementary materials. Holm-corrected post-hoc tests conducted for each of these variables revealed significant differences between partial responses from action cancellations and those from changes of mind (in both the standard and infrequent flanker tasks). Specifically, action cancellation partial responses had greater amplitude, length, and occurred at shorter latency than change of mind partial responses. With regards to the model conducted on offset slopes, there was a main effect of condition and congruence, but no interaction effect.

## 4. Discussion

Over two experiments featuring novel flanker tasks and stop signal tasks (SSTs), we assessed the generalisability of the behavioural slowing and suppression of muscle activity that has been proposed to occur in response to salient, infrequent stimuli (Diesburg & Wessel, 2021). We postulated, based on this framework, that in a flanker task where flanking arrows occurred infrequently (33% of trials), there would be a generic slowing relative to a conventional flanker task, regardless of the congruency of the flankers. Contrary to this prediction, we observed that only infrequent incongruent flankers resulted in behavioural slowing (relative to a standard flanker condition) and only after four preceding trials without any flanker arrows. We also predicted that presenting flanker stimuli infrequently *and* after a delay would result in a pause process observable at the muscle as a partial response, occurring with a latency that was not significantly different to that of partial responses in stopping tasks. Despite the fact that partial responses occurred, as expected, following the presentation of stop signals in SSTs in a significant proportion of trials, their occurrence was extremely rare following delayed infrequent flankers. While this precluded a planned analysis of the latency of these partial bursts, it strongly suggests that presenting an unexpected stimulus after the initiation of an action does not elicit a generic and automatic pause process, challenging prominent theoretical models. Instead, in the context of SSTs, partial responses likely reflect processes that are specific to action cancellation, challenging two-stage models which have predominantly attributed them to pause processes associated with attentional capture across multiple task contexts, based on the rapid latency with which they occur (Diesburg & Wessel, 2021; Tatz et al., 2021; Wadsley et al., 2023).

### 4.1 Behavioural slowing to infrequent and salient stimuli is highly context sensitive

Presenting flankers infrequently only resulted in slowed reaction times (relative to the Standard Flanker Task) under very specific conditions: when flankers were incongruent with the central stimulus, and when they occurred following at least four consecutive trials *without* flankers (figure 3A). A growing body evidence suggests that activation of the rIFG and involuntary behavioural slowing occurs to infrequent and unexpected stimuli (Sebastian et al., 2021; Wessel & Aron, 2017). Past research has argued that stop signals in stop signal tasks (occurring on 25-33% of trials) are sufficiently infrequent and unexpected to elicit this response, and that this suppression is distinct from voluntary action cancellation (Diesburg & Wessel, 2021). Support for this claim primarily comes from *stimulus selective SSTs,* which include “ignore” signals. Ignore signals are equivalent in salience and frequency to stop signals, but do not require the cancellation of movement (Bissett & Logan, 2014; van de Laar et al., 2011; Weber et al., 2026). Past research has observed behavioural delays, partial responses, suppression of corticospinal excitability, and electrophysiological signatures of stopping to ignore stimuli - and some authors have interpreted this as evidence of a generic pause process occurring (Kemp et al., 2024; Tatz et al., 2021; Wadsley et al., 2023; Weber et al., 2024). However, the interpretation that these occur on account of attentional capture and will generalise to other contexts (Diesburg & Wessel, 2021) isn’t parsimonious if, in those other tasks (where stopping is *not* expected) equivalently infrequent salient stimuli don’t trigger these effects. Instead, it seems more plausible that when participants need to discriminate between numerous possible stimuli to determine if stopping is required, they adopt a strategy of stopping to *all* stimuli, and then resume movement when it is deemed appropriate (Bissett & Logan, 2014; Sanchez-Carmona et al., 2021). As such, many observations (behavioural, neural, electrophysiological) to ignore stimuli may simply reflect a transient enactment of a single stopping network (rather than a distinct “pause” mechanism), that is rapidly released before movement is reinitiated. Or, given dynamic and rapid updating of motor plans immediately following action cancellation is a known capacity of the nervous system (Gronau et al., 2024), an additional, separate movement may be enacted in response to the ignore signal.

If we assume a pause process did influence performance in the Infrequent Flanker Task, it must – on account of only being behaviourally manifested in incongruent trials – occur after some degree of stimulus feature decoding. This doesn’t align with the theoretical model, which describes a generic and rapid process (Diesburg & Wessel, 2021), but more importantly, overlooks other, more parsimonious explanations. For example, broader adjustments to perceptual inhibition may be implemented when participants are actively filtering out peripheral stimuli, but which lapse (perhaps as an energy/cognitive-resource saving strategy) when they are no longer needed. Put differently, when perceptual systems have adapted to solely responding to an unambiguous central stimulus (with no flankers) for a number of trials in a row, the cognitive resources allocated to perceptual inhibition may be downregulated. Cognitive modelling has described the interference from flankers as being subject to a shrinking spotlight of attention, whereby flanking stimuli compete with processing of the central stimulus to a greater degree *early* in the trial, and this influence gradually reduces over the course of a single trial (White et al., 2011). The processes determining the size and rate of change of this spotlight may be subject to sequence effects over numerous trials, reflecting the balance of participants’ cognitive effort and energy saving strategies. This could account for the greater slowing to incongruent trials specifically after at least four trials without flankers. This argument could also be framed in terms of conflict monitoring accounts, which posit that conflict triggers increased allocation of cognitive resources (Botvinick et al., 2004; Clayson & Larson, 2011): In the absence of any flanking stimuli that need to be ignored, cognitive resources may be progressively reduced, such that once an incongruent trial is presented, participants are more susceptible to the influence of conflicting information. Ultimately, these interpretations seem more parsimonious than a pause process that is sensitive to both stimulus features and sequence effects.

#### 4.2 Partial responses to infrequent stimuli only manifest under specific task conditions

It may be that a pause mechanism only manifests in SSTs because the infrequent signal is presented *after* the go signal, once a movement is selected and initiated. To test whether this pause generalises to a task that does not involve action cancellation, we devised a Delayed Infrequent Flanker Task, where flankers appeared after the go stimulus, commensurate with the stop signal presentation in SSTs. Contrary to the pause then cancel model predictions, the absence of both partial responses (relative to stopping tasks) and additional behavioural slowing (relative to the Standard Flanker Task) to infrequent stimuli (see Table 5) suggests that salient infrequent stimuli do not reliably elicit a pause outside of stopping contexts.

While flanker stimuli certainly influence behavioural performance, one could argue that the infrequent flankers did not trigger the pause because they were not “task relevant” (participants were instructed to ignore them). But the same is true of “ignore” stimuli in stimulus selective SSTs, and these have been interpreted as supportive evidence *for* the pause then cancel model (Tatz et al., 2021; Wadsley et al., 2023; Weber et al., 2023). Furthermore, we observed in the current experiment that *infrequent* incongruent flanker stimuli resulted in greater slowing of responses. Given this influence on behaviour, it seems plausible that presenting these stimuli infrequently and after a delay would trigger a pause process, if such a process generalises beyond stopping contexts. Despite an almost complete absence of partial responses in the Delayed Infrequent Flanker Task precluding our planned analysis (comparison of latencies between this and stopping tasks), we believe this highlights a key limitation of the pause then cancel model, whereby it currently does not explain the role of proactive adjustments to the motor system that occur in anticipation of stopping (Kenemans, 2015) – manifesting behaviourally in the current task as proactive slowing (see Figure 4). If a two-stage model of action stopping is correct, perhaps these proactive adjustments are necessary for signatures of the pause process to manifest.

### 4.3 Variable stopping speed across conditions demonstrates sensitivity to stop signal salience

Following filtering of slow CancelTimes which were unlikely to represent rapid inhibitory processes (>200ms; Figure 6), stopping was significantly faster in the Stop With Flankers Task than the Standard SST. This corroborates findings that action cancellation can be facilitated when stop signals are more salient, either as a result of recruiting greater low-level sensory resources (on account of taking up more space on the screen) or minimising potential refractoriness of sensory processing that occurs if stop signals are presented in the same location as go signals (Friehs et al., 2024; Kenemans, 2025; Weber et al., 2026). It has been argued that, if partial responses do not solely represent a pause, they may instead reflect a *combination* of the pause *and* cancel processes, whereby faster stops are predominantly underpinned by the pause mechanism, and slower stops by the cancel mechanism (Hervault & Wessel, 2025). This explanation could be viewed as parsimonious with the current result where the more salient stimuli in the Stop With Flankers Task recruited the pause mechanism more regularly, and this resulted in faster CancelTimes. However, our previous work considering partial response distributions across multiple datasets does not provide weight to the notion that action cancellation relies on two distinct events with different temporal profiles (Salomoni et al., 2025). Nonetheless, the current result suggests that imaging research (for instance, with fNIRS) coupled with electromyography, specifically comparing trials with fast and slow stops, could provide further insights. For example, results might support a two-stage model of stopping by demonstrating greater activation of the rIFG (purported origin point of the pause process) in faster stops, and greater activation of the preSMA (purported origin point of the cancel process) in slower stops (with stopping speed indexed on a trial-by-trial basis via EMG partial responses).

The number of go trials which preceded the stop signal had no significant influence on physiological indicators of stopping speed, suggesting the mechanism underpinning reactive stopping is resistant to sequence effects. Recent work has demonstrated that SSRT is faster when stop signals are presented more frequently (20% of trials vs 50%) (Doekemeijer et al., 2023). This was interpreted as the result of broader *proactive* adjustments that prime the motor system for reactive stopping. The authors ran an additional analysis to test for potential sequence effects by calculating SSRT exclusively based on go and stop trials that had been preceded by a go trial, with the main result replicated. In the current experiment, the use of single-trial indices allowed for a more robust test of potential sequence effects on stopping speed. While proactive mechanisms can influence the implementation of stopping, our results also support the claim that these reflect overarching strategic adjustments, rather than trial-to-trial fluctuations in expectation based on recency. Notably, this contrasts with the high contextual specificity and sequence effects we observed in the Infrequent Flanker Task, whereby slowing only occurred to incongruent trials, and only after four non-flanker trials. Again, this suggests that distinct mechanisms underpin responses to salient, infrequent stimuli depending on context (rather than muscle suppression in SSTs representing a generalisable suppression that is specifically due to attentional capture).

#### 4.4 Electrophysiological differences between changes of mind and action cancellation

Our exploratory analysis comparing partial responses associated with a “change of mind”, and those associated with action cancellation revealed a number of differences, though most are intuitive to understand as a product of the tasks used to elicited them. For instance, the greater amplitude and length of partial responses following stop signals likely reflect the fact that these responses have more time to develop before suppression occurs: In SSTs participants are constantly on the brink of failing to stop as a result of the SSD staircasing algorithm (∼50% of initiated responses cannot be cancelled effectively), resulting in a large number of trials where the amplitude of the cancelled movement was almost at the threshold where a button press would be registered. In contrast, flanker tasks demonstrate relatively few trials where participants fail to change their mind before pressing the wrong button (so fewer partial responses approach the threshold that would result in an incorrect button press). We did also observe differences in latency between these processes, relative to the signals that they are in response to, whereby changes of mind occurred more slowly than action cancellation. If changes of mind represent the outcome of gradually accumulating evidence from competing response options, they are subject to the speed of this decision-making process (Servant et al., 2015). In contrast, the required response to a stop signal is unambiguous and features no decision-making process based on stimulus features (unless stimulus selective stopping tasks are used; Weber et al., 2026), resulting in earlier muscle suppression, as such this observation reflects the differing cognitive processes that precede muscle suppression in these contexts. The latency of partial responses associated with changes of mind was also influenced by congruency (Figure 5c). This aligns with previous research observing faster changes of mind to congruent (213ms) than incongruent (231ms) trials (Servant et al., 2015), but also extends this to show that these differences reflect facilitation and interference (respectively) relative to neutral trials. Cognitive modelling has captured the rate and latency of partial responses in this context by describing them as the outcome of evidence accumulation processes which, once crossing a threshold, begin to recruit muscle fibres. This activation remains subject to evidence accumulation processes (until a second threshold, representing the behavioural response, is crossed) enabling a change to another response should the weight of the evidence change (Servant et al., 2015). Extending similar cognitive modelling approaches to compare and interpret partial responses in the context of action cancellation likely represents a worthwhile avenue for future research.

### 4.5 Conclusion

If the pause then cancel model of stopping accurately describes human behaviour, the short latency of partial responses following stop signals requires that they be, at least in part, attributed to the pause process. This process is purportedly triggered by attentional capture and is not specific to contexts where stopping may be required. The current findings challenge this description, showing that fast, automatic inhibition fails to generalise to equivalently infrequent stimuli outside action cancellation contexts. Given that a pause seems to only reliably manifests at the muscle in contexts where stopping is required, the functional role of such a process, and the contexts in which it would be generated, requires further research/specification.

As a direction for future research, we highlight that the suppression of movement is common to both changes of mind and action cancellation, though one occurs in response to internal decision-making processes, while the other occurs in response to an external stimulus. Plausibly, overlapping mechanisms are common to the cancellation of movement in both contexts, though no work to our knowledge has investigated potential shared neural correlates. Instead, the attribution of specific neural mechanisms to partial responses, based on their latency and context, has thus far been driven by theoretical accounts. The current result challenges a prominent account, and highlights how further empirical support is required to inform current theory. Coupling similar EMG analysis (to isolate trials with action cancellation) with compatible brain imaging (e.g., fNIRS) and/or brain stimulation techniques, will likely provide an avenue to further specify frameworks for understanding of human executive control and decision-making.

## Data Statement

The data and analysis script for this manuscript can be found here: https://osf.io/5tnx8 Our MATLAB scripts for analysing EMG data can also be found on OSF: https://osf.io/r8a54

## Credit Statement

Simon Weber – Conceptualisation, Methodology, Formal Analysis, Software, Writing - Original Draft Preparation

Keegan Haugh - Conceptualisation, Investigation, Data Curation, Writing - Original Draft Preparation

Sauro E. Salomoni – Conceptualisation, Formal Analysis, Methodology, Software, Writing - Review & Editing

Ahreum Lee - Conceptualisation, Investigation, Data Curation, Writing - Original Draft Preparation

Evan J. Livesey - Conceptualisation, Writing - Review & Editing

Mark R. Hinder - Conceptualisation, Supervision, Writing - Review & Editing

## Supplementary Materials

### Comparisons of Partial Response Attributes

#### Amplitude

The model comparing amplitudes of partial responses revealed a main effect of condition *F*(2,2285) = 63.78, *p* < .001, a main effect of congruence *F*(2,2767) = 16.43, *p* < .001, and a significant interaction effect *F*(4,2759) = 4.81, *p* < .001. Holm corrected post-hocs were conducted checking the effects of condition at each level of congruence. For congruent trials, action cancellation partial responses demonstrated significantly greater amplitude than change of mind partials in both the standard, *t*(2711)= 5.059, *p* <0.001, and infrequent, *t*(2780) = 7.359, *p* <0.001, flanker conditions, though no diffrence was observed between infrequent and standard flanker conditions *t*(2780) = 1.36, *p* = 0.176. For incongruent trials, results were similar: Action cancellation partial responses had greater amplitude than change of mind partial responses in standard, *t*(2696)= 6.26, *p* < 0.001, and infrequent, *t*(2639) = 3.68, *p* <0.001, flanker conditions, though there was also a significant diffrence between infrequent and standard flanker conditions *t*(2767) = 2.91, *p* = 0.003. In neutral trials, action cancellation partial responses had greater amplitude than change of mind partials in standard, *t*(2689)= 6.67, *p* < 0.001, and infrequent, *t*(2750) = 7.06, *p* <0.001, flanker conditions, though there was no significant diffrence between infrequent and standard flanker conditions *t*(2771) = 0.15, *p* = 0.879.

#### Latency

The model run comparing latencies of partial responses revealed a main effect of condition, χ^2^(2) = 997.25, *p* < 0.001, a main effect of congruence, χ^2^(2) = 57.55, *p* < 0.001, and a significant interaction effect, χ^2^(4) = 14.04, *p* < 0.001. Holm corrected post-hocs revealed that for congruent trials, action cancellation partial responses were faster than change of mind partials in the standard, *z* = 11.96, *p* <0.001, and infrequent, *z* = 13.10, *p* <0.001, flanker conditions, though no diffrence was observed between infrequent and standard flanker conditions *z* = 0.17, *p* = 0.866. The same pattern was observed for incongruent trials, with faster suppression in action cancellation compared to standard, *z* = 23.18, p <0.001, and infrequent flankers, z = 26.76, p < 0.001, but no difference between the flanker conditions, z = 1.72, p = 0.085. Neutral trials also demonstrated faster suppression in action cancellation compared to change of mind partials in standard, *z* = 14.94, p <0.001, and infrequent flankers, z = 17.90, p < 0.001, and no difference between the flanker conditions, z = 1.40, p = 0.161.

#### Length

The model comparing the length of partial responses revealed a main effect of condition *F*(2,2744) = 43.54, *p* < .001, a main effect of congruence *F*(2,2754) = 12.33, *p* < .001, and a significant interaction effect *F*(4,2750) = 4.64, *p* = 0.001. Holm corrected post-hocs revealed that for congruent trials, action cancellation partial responses had significantly greater length than change of mind partials in the standard, *t*(2788)= 5.11, *p* <0.001, and infrequent, *t*(2782) = 7.652, *p* <0.001, flanker conditions, though no diffrence was observed between infrequent and standard flanker conditions *t*(2759) = 1.53, *p* = 0.126. Similarly, for incongruent trials action cancellation partial responses had significantly greater length than change of mind partials in the standard, *t*(2787)= 2.93, *p* = 0.007, and infrequent, *t*(2785) = 3.592, *p* = 0.001, flanker conditions, though no diffrence was observed between infrequent and standard flanker conditions *t*(2755) = 0.37, *p* = 0.712. In neutral trials, action cancellation partial responses had significantly greater length than change of mind partials in the standard, *t*(2788)= 4.61, *p* < 0.001, and infrequent, *t*(2786) = 5.29, *p* < 0.001, flanker conditions, and again, no diffrence was observed between infrequent and standard flanker conditions *t*(2755) = 0.22, *p* = 0.828.

#### Offset slope

The model run comparing offset slopes revealed a main effect of condition, *F*(2,278) = 3.37, *p* = 0.035, a main effect of congruence, *F*(2,277) = 3.50, *p* = 0.030, and no significant interaction effect, *F*(4,2769) = 0.50, *p* = 0.733. Holm corrected post-hocs investigating the main effect of condition revealed a marginally significant difference between action cancellation partial responses and those in the infrequent flanker task *t*(2788) = 2.38, *p* = 0.052. The comparison between the action cancellation responses and those in the standard flanker task was not significant, *t*(2768)= 2.04, p = 0.083, nor was the comparison between the infrequent and standard flanker tasks *t*(2781)= 0.07, p = 0.943. The main effect of congruence was underpinned by a significant difference between incongruent and neutral trials *t*(2770) = 2.48, *p* = 0.040, though there was no difference between incongruent and congruent trials, *t*(2773) = 0.03, *p* = 0.974, nor congruent and neutral trials *t*(2773) = 2.10, *p* = 0.072.

While we did not formulate a formal hypothesis, steeper offsets for action cancellation may have been predicted on account of specialised inhibitory mechanisms driving this suppression. Alternatively, we note that a change of mind partial response is followed by rapid initiation of a response on the other side. Initiation of a unilateral movement triggers interhemispheric inhibition, which serves to suppress the homologous muscle on the contralateral limb (Hinder et al., 2010). As such comparison of offset slopes in SSTs and stop change tasks (which require cancellation and the subsequent enactment of a response on the other side), may warrant further investigation. Currently, a key challenge in the inhibition literature is that purported neural mechanisms, which often suggest a two-stage model, are not reconcilable with observations at the level of the muscle, which instead suggest a single stopping mechanism (Chaudhuri et al., 2025; Salomoni et al., 2025). Finding novel ways to link signatures at the muscle (with parameters beyond presence/absence of partial burst; Kemp et al., 2024) with neural signatures, may help to clarify this debate.

## Supplementary CancelTime Results

**Figure Sup1:**
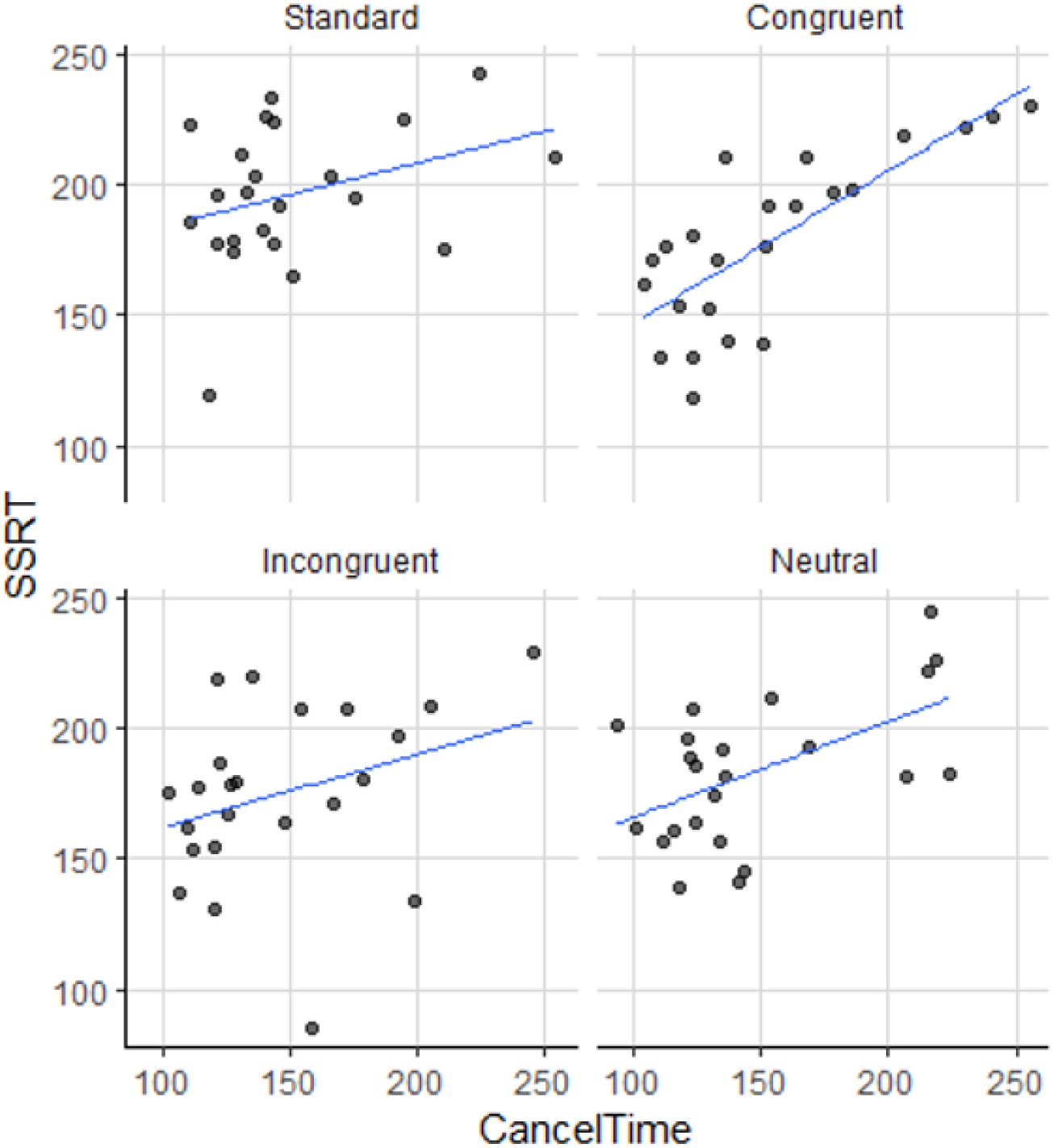
Correlations of participant mean canceltimes with integration method calculated SSRTs for each stop type. *Interpretation of CancelTime distributions split by SSD*

**Figure Sup 2:**
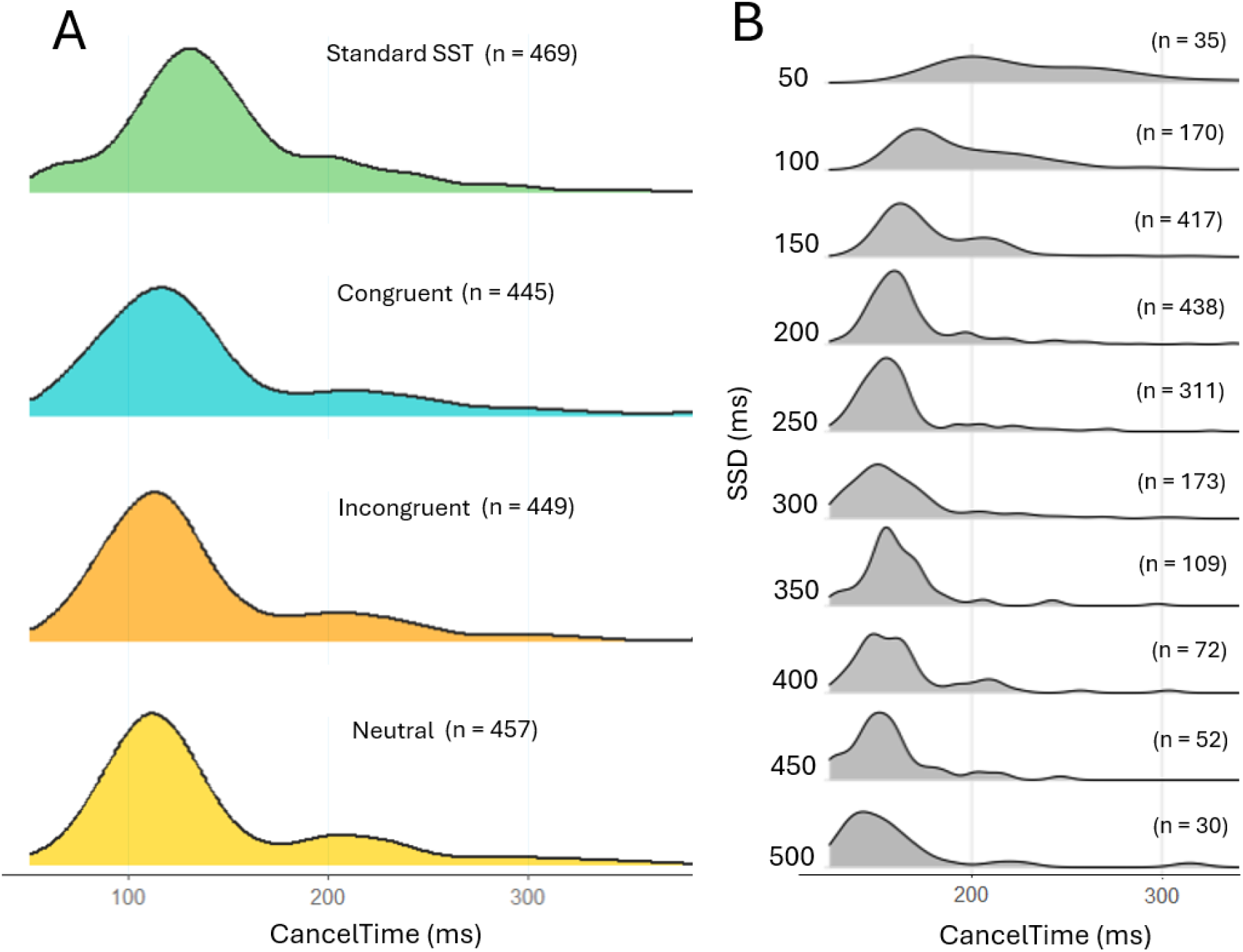
**A)** Canceltime distributions from the standard SST, and each level of congruence from the Stop With Flankers task. The model revealed no significant differences between these conditions. **B)** Interpreting the shape of CancelTime distributions becomes clearer when SSD effects are considered, with flatter, more variable distributions occuring at low SSDs (data is pooled across each type of stop signal). Note that there were also 6 trials at an SSD of 550ms, and 7 tria at an SSD of 600ms (not represented).

